# Regulation of Type VI secretion systems of *Burkholderia thailandensis* during transition to intracellular lifestyle

**DOI:** 10.64898/2026.01.30.702787

**Authors:** Hoi Ching Cheung, Daria Jacqueline Wüst, Miro Thorsten Wilhelm Plum, Patricia Reist Iscar, Marek Basler

**Affiliations:** Biozentrum, University of Basel, Spitalstrasse 41, CH-4056 Basel, Switzerland

## Abstract

Pathogenic bacteria tightly regulate gene expression in response to environmental cues. In *Burkholderia* species, the Type VI Secretion Systems (T6SS) mediate interbacterial competition and host cell interactions. Here, we show that *B. thailandensis* switches from the antibacterial T6SS-1 to the anti-eukaryotic T6SS-5 during phagosomal maturation. While T6SS-1 expression persists in the host cells, its assembly progressively declines with increasing T6SS-5 expression. This switch is controlled by the two-component system VirAG as deletion of *virA* blocks T6SS-5 expression and restores T6SS-1 assembly in host cells. Notably, T6SS-1 activity in host cells induces mitochondrial fragmentation and apoptosis. Our data suggest that *Burkholderia* has evolved a mechanism to silence T6SS-1 upon host entry to preserve its replicative niche.

## Introduction

To adapt to various niches, pathogens employ different factors required for virulence and survival in the environment. The expression of these factors is often tightly controlled. The facultative intracellular pathogen *Burkholderia pseudomallei* is endemic in tropical and subtropical regions and is the causative agent of the disease Meliodosis^1–4^. Patients are commonly infected via inhalation, ingestion or skin abrasions^5^. It is estimated to have 165,000 meliodosis cases leading to 89,000 deaths per year globally^6,7^, and the mortality rate ranges from 10% to 40%^4,8,9^. Due to the disease’s high mortality rate and resistance to antibiotics, *B. pseudomallei* is listed as a tier 1 select agent by the US Centers for Disease Control and Prevention^10,11^. The closely related, less virulent *Burkholderia thailandensis* is often used as an attenuated model for studying *Burkholderia* virulence^12^.

Both *B. pseudomallei* and *B. thailandensis* are able to infect a range of phagocytic and non-phagocytic eukaryotic cells^13^. Upon entry into host cells, *B. thailandensis* escapes the phagosome using effectors of the Type III Secretion System (T3SS_Bsa_) ^14^. *B. thailandensis* moves in the host cytosol by using an intracellular flagellum and by polymerising host cell actin with the bacterial protein BimA, which is a mimic of the eukaryotic Arp2/3 complex^15^. Actin-tails promote formation of membrane protrusions into the neighbouring host cell. In the protrusions, *B. thailandensis* assembles its T6SS-5 to fuse host cell membranes by VgrG-5, leading to a protrusion escape or fusion of host cells^16–20^.

T6SS is structurally homologous to a contractile bacteriophage tail and functions by propelling a needle-like structure with effectors into a neighbouring target cell^21–25^. Its core structure consists of a membrane and baseplate complex formed around a VgrG-PAAR spike^26–32^. The sheath is formed by the assembly of TssB-C around an inner tube of Hcp rings starting from the VgrG-PAAR base and extending into the cytoplasm. When the TssB-C sheath contracts, it drives the Hcp-VgrG-PAAR and associated effectors outside the cell and into the target cell^33,34,35,36^. The contracted sheath is disassembled by the ATPase ClpV^25,37–39^.

*B. pseudomallei* and *B. thailandensis* have multiple T6SS with distinct roles and high homology between the two species^16,40^. T6SS-5 was shown to be important for cell-to-cell spread^17,18,20,41,42^; whereas T6SS-1 is required for bacterial killing^16,42,43^. In previous work, functional chromosomal fusion of the sheath protein TssB of both T6SS-1 and T6SS-5 with fluorophores have been generated^18,44^.

The expression and assembly *B. thailandensis* T6SS-5 are tightly regulated. The regulatory cascade starts with BsaN, a central regulator located in the T3SS_Bsa_ locus. It regulates T3SS effectors and two downstream T6SS-5 regulators VirAG and BprC^45,46^. A two-component histidine kinase, VirAG, regulates T6SS-5 expression during infection; and BprC regulates T6SS-5 expression when grown in medium, except for the *tssBC* operon^45^. The sensor VirA phosphorylates VirG upon sensing host cytosol when the bacterium escapes from the endosome and the phosphorylated VirG binds to the promoter regions of TssB-5 and Hcp-5^47^. However, the signals for BsaN and BprC activation are unknown and the redox signal sensed by VirAG is the only signal identified during infection. The regulation of the anti-bacterial T6SS-1 is poorly understood in *B. thailandensis*. It was shown that quorum sensing controls the expression of T6SS-1 effectors and VgrG-1^43^. In terms of post-translational regulation of T6SS-1, it was shown that T6SS accessory protein TagM-1 is important for T6SS-1 assemblies^44^.

In this study, we aim to understand how *Burkholderia* regulates its different T6SS in the environment and during infection. We investigated the expression and assembly of all five T6SS of *B. thailandensis* by live-cell imaging in medium and during infection. We observed that the anti-eukaryotic T6SS-5 is expressed before the bacterium escapes to the host cytosol, and that the anti-bacterial T6SS-1 assemblies are inhibited during infection. The phagosomal environment is a transition state where there is gradual increase in T6SS-5 expression and diminishing T6SS-1 assemblies. Furthermore, we identified VirAG as a regulator for both T6SS-5 expression and T6SS-1 assemblies. Deletion of *virA* abolished T6SS-5 expression and allowed T6SS-1 assemblies during infection, which led to apoptotic host cell death. This indicates that switch between the two T6SS allows the bacteria to adapt to their environment and to spread during infection.

## Results

### T6SS-2, T6SS-4 and T6SS-6 are not expressed during infection of A549 cells

*B. thailandensis* harbour five T6SS on different chromosomal loci and with different functions. Previous studies have mainly focused on the anti-eukaryotic T6SS-5 as it alone was sufficient to cause virulence in mice^16^. A transcriptomics study has hinted that T6SS-2, T6SS-4, and T6SS-6 have increased expression when inside of host cells^48^. However, it remains unclear if these T6SS are expressed intracellularly. To test this, we constructed strains of *B. thailandensis* with the T6SS sheath proteins TssB tagged with mNeonGreen. We then infected a monolayer A549 cells and performed live-cell imaging. Intracellular bacteria were identified by either expression of mCherry2 under a ribosomal promoter, or expression of TssB-5-mCherry2. Images were acquired between 11-12 hours post-infection (hpi). We acquired images of bacteria in the cytoplasm and in membrane protrusions. We found that the T6SS clusters T6SS-2, T6SS-4 and T6SS-6 were not expressed during infection (Figure S1A). In accordance with previous studies, the T6SS-5 was expressed in *B. thailandensis* at 12 hpi. Notably, the anti-bacterial T6SS-1 expression was detected during infection (Figure S1B).

We next assessed T6SS expression when grown in liquid culture and imaged on LB (Luria–Bertani) agar pads. T6SS-2 and T6SS-4 are not expressed under such conditions as the fluorescently-labelled *B. thailandensis* strains exhibited background fluorescence levels comparable to those of the unlabelled strain. The T6SS-6–labelled strain exhibited heterogeneous expression, but no assemblies were detected (Figure S1C).

### T6SS-5 is repressed *in vitro* and induced within host cells before phagosomal escape

The anti-eukaryotic T6SS-5 was shown to be expressed when *Burkholderia thailandensis* locates in the host cell cytosol, however, T6SS-5 expression dynamics has never been analyzed^47^. In accordance with literature, we detected no expression of T6SS-5 when *B. thailandensis tssB5-mScarlet-I* was imaged *in vitro* on agar pads while and its expression was induced by expressing VirAG from an arabinose-inducible plasmid (Fig S1D).

To investigate the T6SS-5 expression dynamics during infection, we infected a monolayer of A549 cells with *B. thailandensis tssB5-mScarlet-I* constitutively expressing cytosolic mCherry2 under a ribosomal promoter pSC12 at a Tn7 insertion site. Host cell membranes were stained with CellMask deep red for 5 min and washed before imaging. Kanamycin was added to the medium to kill extracellular bacteria. Images were taken at 1-minute intervals to identify the time point when TssB-5-msfGFP is expressed. During a 5-hour imaging period, we detected a signal from TssB-5-msfGFP at 3 hours 10 minutes post-infection (Fig 1B, Video 1). Furthermore, we tracked the bacteria and measured change in TssB-5-msfGFP fluorescence intensity in time. A region free of host cells and bacteria was selected at each timepoint to measure background green fluorescence. By performing a two-way ANOVA with Tukey’s multiple comparisons test, we identified the time-point where T6SS-5 expression is significantly different from the background signal (Fig 1C). Doing similar analysis on 116 bacteria across three biological replicates, we observed that most bacteria express T6SS-5 between 2 to 4 hpi (Figure 1F). Furthermore, we imaged T6SS-5-expressing bacteria at 10-30 seconds intervals and found that *B. thailandensis* assembled T6SS-5 when inside of the phagosome (Fig 1G, S2). Notably, the T6SS-5 assemblies did not disrupt the phagosomal membrane. This contrasts with later stages, when T6SS-5 assemblies is known to facilitate lyse host cell membranes for cell-to-cell spread^18,19^.

**Figure 1.**
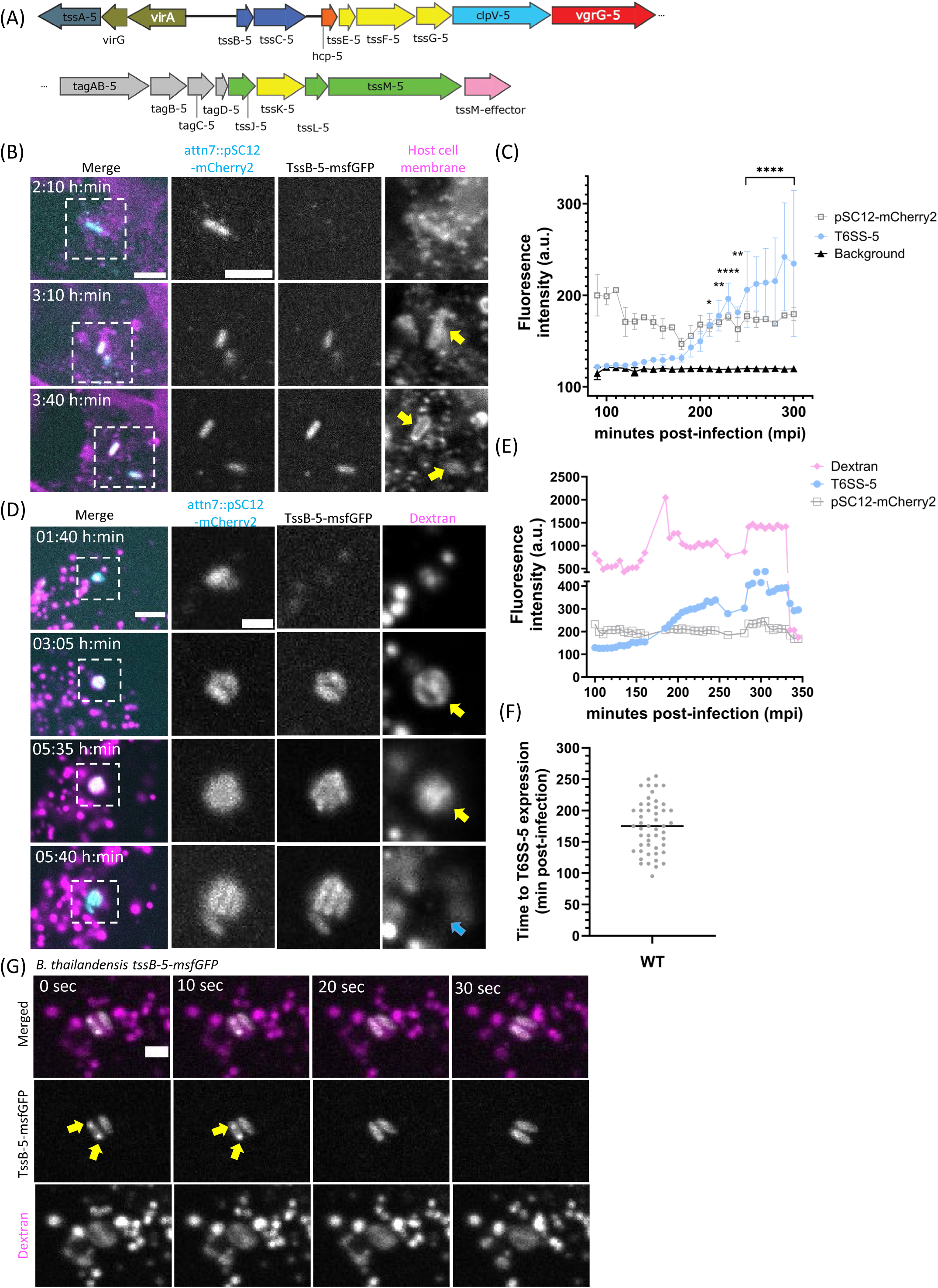
Expression and assembly of T6SS-5 and its regulatory component VirAG inside primary vacuole. **(A)** Scheme of the anti-eukaryotic T6SS-5 genetic cluster. **(B)** Time-lapse imaging of A549 cells infected with *B. thailandensis* expressing cytosolic mCherry2 under ribosomal promoter pSC12 (cyan) and TssB-5-msfGFP (grey) at MOI of 50. Host cell membrane was stained with CellMask DeepRed (magenta). Yellow arrows indicate host cell membrane around the bacteria. Scale bars represent 5 µm. Images are representative of at least 3 biological replicates. **(C)** Fluorescence intensity of various channels during the 5-hour infection period shown in (B). Bacteria were segmented and tracked, and fluorescence intensities of TssB-5-msfGFP (T6SS-5, in blue), bacterial cytosol (pSC12-mCherry2, in grey), and background (black) were plotted. A two-way ANOVA with Tukey’s multiple comparisons test was performed (* P<0.05, ** P<0.01, *** P<0.0001). **(D)** Time-lapse imaging of A549 cells infected with *B. thailandensis* expressing cytosolic mCherry2 under ribosomal promoter pSC12 (cyan) and TssB-5-msfGFP (grey) at MOI of 200. Host cells were pre-treated with 0.25 mg/mL of fluorescent dextran for 18 hours (magenta). Yellow arrows indicate two bacteria expressing T6SS-5 when inside of a dextran-labelled vacuole. Blue arrow indicates the disappearance of the dextran signal. Scale bars represent 5 µm in the overview image and 5 µm in the magnified image. Images are representative of at least 3 biological replicates. **(E)** Fluorescence intensity of various channels during the 5 hour 45 minutes infection period shown in (D). Bacteria were segmented and tracked, and fluorescence intensities of TssB-5-msfGFP (T6SS-5, in blue), bacterial cytosol (pSC12-mCherry2, in grey), and fluorescent dexran (magenta) were plotted. **(F)** Scatter plot showing the timing of T6SS-5 expression. Each dot represents an individual bacterium. A549 cells were infected with *B. thailandensis* expressing TssB-5-mScarlet-I at MOI 50-200. Time-lapse mages were acquired from 1:30 hpi at either 1 min, 5min, or 10 min intervals. **(G)** *B. thailandensis* assembling T6SS-5 at 3 hpi inside of a dextran-labelled vacuole during infection of A549 cells. A549 cells were infected with *B. thailandensis* expressing TssB-5-msfGFP (grey) at MOI 200. Host cells were pre-treated with 0.25 mg/mL of fluorescent dextran for 18 hours (magenta). Yellow arrows indicate T6SS-5 assembly events. Scale bar represents 2 µm. Images are representative of at least 3 biological replicates.

Since previous studies focused on the expression of Hcp-5 but not TssB-5^45,47^, we investigated if Hcp-5 was also expressed within our identified time frame of TssB-5 expression. We performed immunofluorescence staining with an anti-Hcp-5 antibody on infected cells fixed at 3 hpi (Fig S3A). As a positive control, a *B. thailandensis* strain expressing VirAG from a pmlBAD plasmid was included, and Hcp-5 was detected. The negative control Δ*hcp-5* strain showed no Hcp-5 expression and for the wild-type, both Hcp-5 and TssB-5 were detected.

Interestingly, we noticed that the bacteria were in a membrane-bound compartment when expressing T6SS-5 (Fig 1B, S3B, yellow arrows). We therefore asked if T6SS-5 is expressed before breakage of the phagosomal membrane. To test this, we pre-loaded A549 cells with 0.25 mg/mL fluorescent dextran 100 kDa overnight, infected the cells with *B. thailandensis* in the absence of dextran, washed with Optimem^TM^ with kanamycin (300µg/mL) after 1 hpi and imaged immediately. We found that *B. thailandensis* expressed TssB-5-msfGFP when co-localized with dextran-labelled vacuoles (Fig 1D, S3D, Video 2). We then tracked the bacteria and measured the fluorescence intensity of TssB-5-msfGFP throughout the infection time course. Similar to Fig. 1C, fluorescence intensity of the dextran, T6SS-5 and bacterial cytosol (pSC12-mCherry2) were plotted (Fig 1E, S3E). The dextran signal remained between 500-2,000 a.u. and only dropped to 200 a.u. after two hours of increased T6SS-5 expression. By analyzing 76 bacteria in dextran-treated A549 cells across three biological replicates, we found that 87% of the bacteria expressed T6SS-5 when colocalized with fluorescent dextran.

### Phagosomal maturation is important for T6SS-5 expression

To elucidate at which step of the phagosomal maturation pathway is *B. thailandensis* expressing T6SS-5, we infected two HeLa cell lines stably expressing Rab5-mApple or Rab7-mApple, and stained with anti-LAMP1 (lysosome-associated membrane protein-1) antibody. We found that most bacteria express T6SS-5 in late phagosomal maturation stages, indicated by colocalization with Rab7-mApple, and that T6SS-5 expressing bacteria often co-localize with LAMP1-positive vacuoles between 2 to 3 hpi (Figure 2B). We quantified over 400 T6SS-5 expressing bacteria in at least 3 biological replicates, and tested for co-localization with either Rab5, Rab7, and LAMP1. Only 2% of T6SS-5 expressing bacteria co-localize with Rab5-labelled phagosomes; whereas 16.5% of those co-localize with Rab7-labelled phagosomes and 29% with LAMP1-labelled phagosomes (Fig 2C).

**Figure 2.**
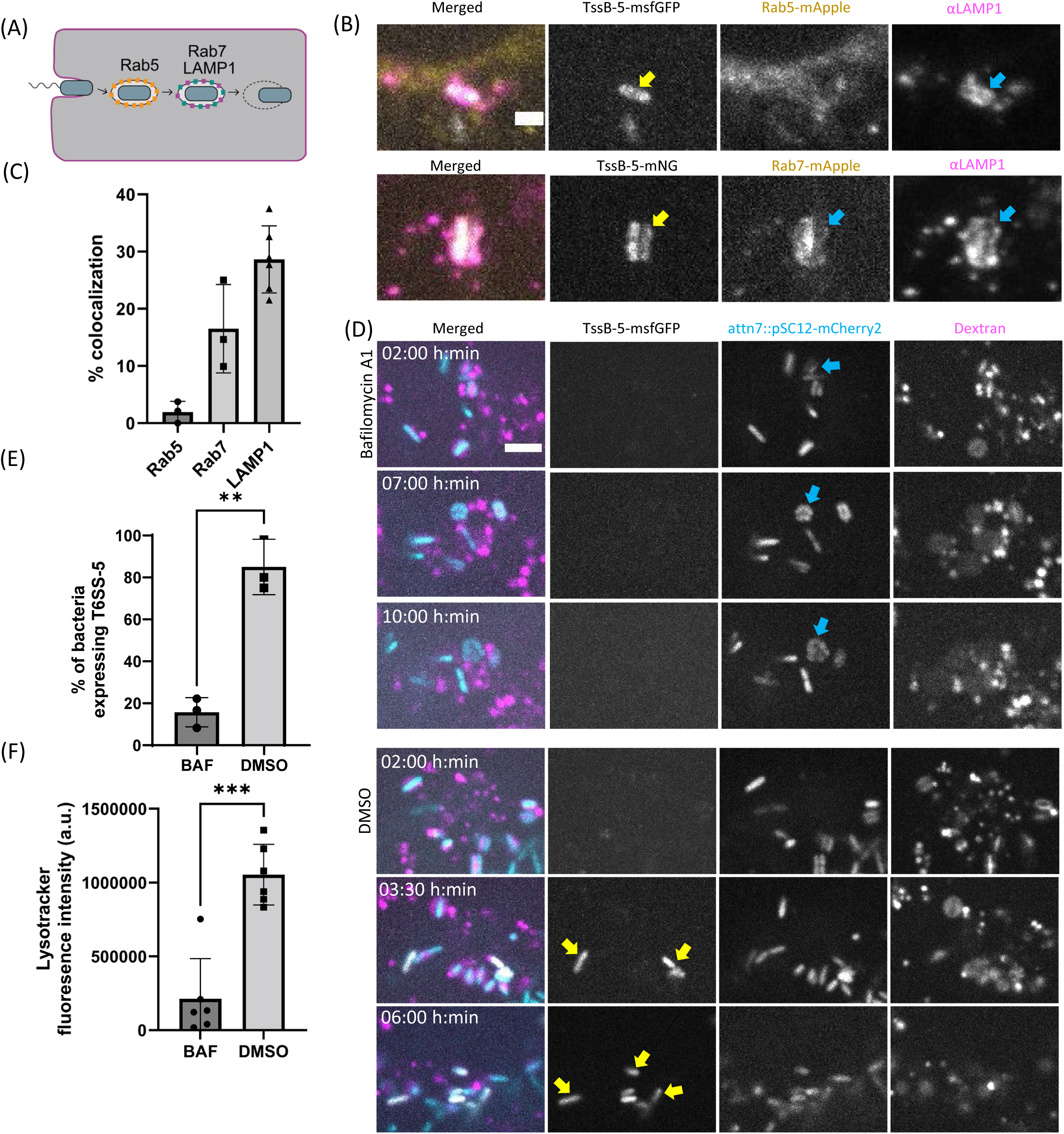
Phagosomal maturation is important for T6SS-5 expression. **(A)** Scheme depicting the phagosomal membrane markers when *B. thailandensis* enters a host cell. **(B)** LAMP-1 immunofluorescence staining of HeLa cells stably expressing Rab5 and Rab7-mApple infected with *B. thailandensis* at MOI 50. Cells were fixed and permeabilized with ice-cold methanol at 3 hpi and stained with anti-LAMP1 antibody at 1 ug/mL for 1 hour at room temperature. Then the slide was washed with PBS, and stained with anti-mouse-AlexaFluor647 overnight at 4 degrees. Yellow arrows indicate TssB-5 expressing bacteria. Blue arrows indicate the presence of phagosomal markers. Scale bar represents 2 µm. **(C)** Quantification of percentage of T6SS-5 expressing bacteria co-localizing with Rab5, Rab7, or LAMP-1 positive vacuoles at 3 hpi. Each data point represents one biological replicate. More than 100 bacteria were counted in each biological replicate. **(D)** Time-lapse imaging of A549 cells treated with bafilomycin A1 (100nM) or DMSO infected with with *B. thailandensis* expressing cytosolic mCherry2 under ribosomal promoter pSC12 (cyan) and TssB-5-msfGFP (grey) at MOI of 200. A549 cells were pre-treated with 0.25 mg/mL of fluorescent dextran for 18 hours to identify intra-phagosomal bacteria. Host cells were pre-treated 1 hour before infection, infected with *B. thailandensis* for 1 hour without treatment, and bafilomycin A1 or DMSO was added and kept in the media with 300 µg/mL kanamycin from 1 hpi onwards. Host cell membrane was stained with CellMask DeepRed (magenta). Blue arrows indicate bacteria that do not express T6SS-1 and yellow arrows indicate T6SS-5 expressing bacteria. Scale bar represents 5 µm. **(E)** Percentage of intracellular *B. thailandensis* expressing T6SS-5 when A549 cells were treated with bafilomycin A1 or DMSO. A549 cells were pre-treated with 0.25 mg/mL of fluorescent dextran for 18 hours to identify intra-phagosomal bacteria. A total of 109 bacteria were identified and tracked for T6SS-5 expression across 3 biological replicates. An unpaired t-test were carried out (** P<0.001). **(F)** Fluorescence intensity of A549 cells treated with either 100nM bafilomycin A1 or DMSO stained with Lysotracker DeepRed. Fluorescent images were adjusted with the same threshold values and integrated density values were measured with ImageJ. An unpaired t-test were carried out (*** P<0.0001).

Since LAMP1 is a common marker for mature, acidified vacuoles^49,50^, we explored the role of phagosomal maturation as a signal for T6SS-5 expression. Acidification of vacuoles can be inhibited by bafilomycin A1^51–53^. We pre-treated A549 cells with 100 nM bafilomycin A1 one hour before infection, infected the cells with *B. thailandensis*, washed with kanamycin at 1 hpi, and replenished the media with bafilomycin A1. At the same time, we also pre-loaded the host cells with 0.25 mg/mL fluorescent dextran 100 kDa overnight to label phagosomes. To ensure bafilomycin A1 inhibits the acidification of vacuoles in our assays, we measured the fluorescence intensity of lysotracker and showed that bafilomycin A1 treatment significantly reduced acidification of vacuoles (Fig 2F). Importantly, the addition of bafilomycin A1 largely abolished T6SS-5 expression (Fig 2D, Video 3). We quantified the percentage of bacteria expressing T6SS-5 with or without bafilomycin A1 treatment (Fig 2E). We analysed 69 bacteria in host cells treated with bafilomycin, and 41 bacteria in host cells treated with DMSO across three biological replicates. Only 8 bacteria (11.5%) expressed T6SS-5 when the host cells were treated with bafilomycin A1, compared to 85% of bacteria expressing T6SS-5 in the DMSO control. Among the 8 T6SS-5-expressing bacteria residing in bafilomycin-treated cells, one bacterial cluster escaped from the phagosome, whereas the rest of the bacteria remained confined within the phagosome (Figure S4A,B). The bacteria were able to replicate inside of dextran-labelled vacuoles but were unable to escape from the phagosome (Fig 2D, Video 3). This indicates that phagosomal maturation and acidification is an important signal for T6SS-5 expression.

### Anti-bacterial T6SS-1 is expressed but not assembled when the bacterium is in the host cell cytosol, and its regulation is dependent on VirAG

It was shown previously that *B. thailandensis* constitutively express T6SS-1 components and they assemble anti-bacterial T6SS-1 when imaged on agar pads^44^. Little is known how they regulate the T6SS-1 during infection. To test this, we infected A549 cells with a *B. thailandensis* strain that expressed the T6SS-1 sheath protein fused to a fluorophore, TssB-1-msfGFP, and performed live-cell imaging after 12 hpi. To our surprise, we observed that the T6SS-1 was expressed but not assembled, as indicated by uniform fluorescence throughout the bacterial cells without the appearance of dynamic puncta typically associated with T6SS assembly events (Fig 3A, S5A, Video 4). To identify the point of T6SS-1 assembly inhibition, we infected A549 cells and acquired time-lapse images of at least 100 different bacteria in unique field of views across different time points. Images were captured every 30 seconds over a 5-minute period to assess whether the bacteria assembled T6SS-1. At early infection time points when the *B. thailandensis* were in dextran-labelled vacuoles, all bacteria expressed T6SS-1, but not all bacteria assembled T6SS-1 (Fig 3B, C). We quantified the percentage of bacteria assembling T6SS-1 at various time points, and found that 33% of intracellular bacteria assembled T6SS-1 between 1-2 hpi. This significantly decreased to 16% when imaged between 2 to 3 hpi. 11% of bacteria assembled T6SS-1 between 3 to 4 hpi and this further decreased to 1% after 4 hpi and 0% after 12 hpi (Fig 3D). In all the time points, none of the cytosolic bacteria assembled T6SS-1. When we infected A549 cells with a *B. thailandensis* strain with both T6SS-1 and T6SS-5 fluorescently labelled, we found that amongst 93 bacteria, when the T6SS-5 is expressed, there are no T6SS-1 assemblies (Fig 3E, S5B). This data suggests that expression of T6SS-5 is negatively correlated with the assembly of T6SS-1.

**Figure 3.**
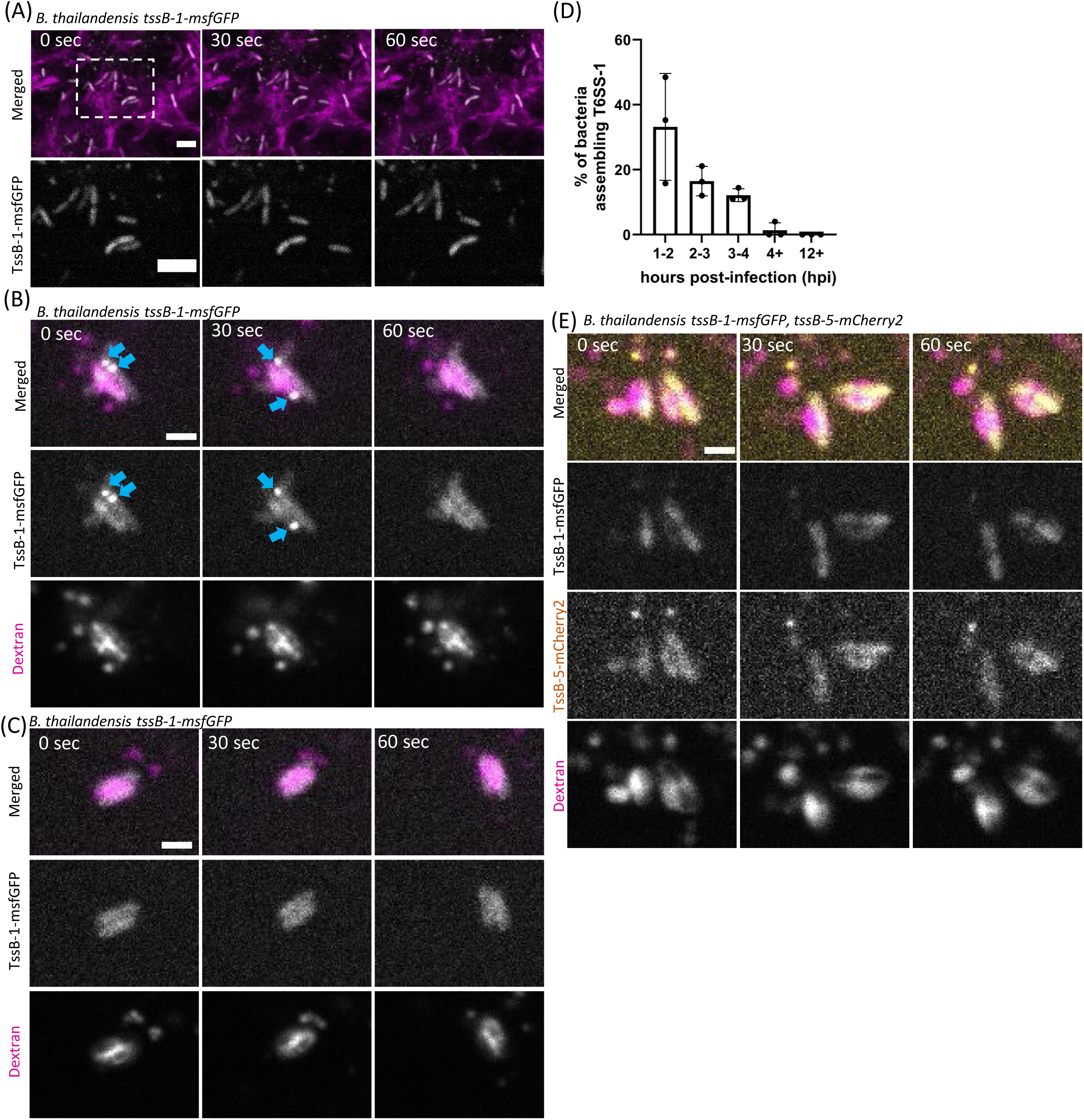
Expression and assembly dynamics of T6SS-1 during infection. **(A)** Time-lapse microscopy of A549 cells infected with *B. thailandensis* TssB-1-msfGFP at MOI of 50 at 13 hpi. Host cell membranes were stained with CellMask DeepRed (magenta). Scale bars represent 5 µm. **(B)** Example of *B. thailandensis* assembling T6SS-1 inside of a dextran-labelled phagosome in A549 cells. Host cells were pre-treated with 0.25 mg/mL of fluorescent dextran for 18 hours (magenta). Images were taken at 45 minutes post-infection. Blue arrows indicate T6SS-1 assembly events. Scale bar represents 2 µm. **(C)** Example of *B. thailandensis* not assembling T6SS-1 inside of a dextran-labelled phagosome in A549 cells. Host cells were pre-treated with 0.25 mg/mL of fluorescent dextran for 18 hours (magenta). Images were taken at 2 hpi. Scale bar represents 2 µm. **(D)** Quantification of the percentage of T6SS-1 assembling bacteria during infection of A549 cells at different time points. A549 cells were infected at an MOI of 200, and at least 100 intracellular bacteria were analyzed per biological replicate (n = 3). Each bacterium was imaged only once in a unique field of view, with time-lapse images acquired every 30 seconds over a 5-minute period. **(E)** Time-lapse microscopy of A549 cells infected with *B. thailandensis* expressing TssB-1-msfGFP and TssB-5-mCherry2 at MOI of 500 at 3 hpi. Host cells were pre-treated with 0.25 mg/mL of fluorescent dextran for 18 hours (magenta). Scale bar represents 2 µm.

### VirAG inhibits anti-bacterial T6SS-1 assemblies during infection

Since VirAG is a two-component system that upregulates the T6SS-5 expression at the beginning of infection, we hypothesized that VirAG also plays a role in inhibiting T6SS-1 assembly. We expressed VirAG from an arabinose-inducible plasmid and imaged T6SS-1 assembly dynamics on agar pads *in vitro* (Fig 4A, Video 5). The parental *B. thailandensis* strain expressing TssB-1-msfGFP assembled T6SS-1 constitutively. However, when VirAG was expressed, T6SS-1 assembly was completely abolished.

**Figure 4.**
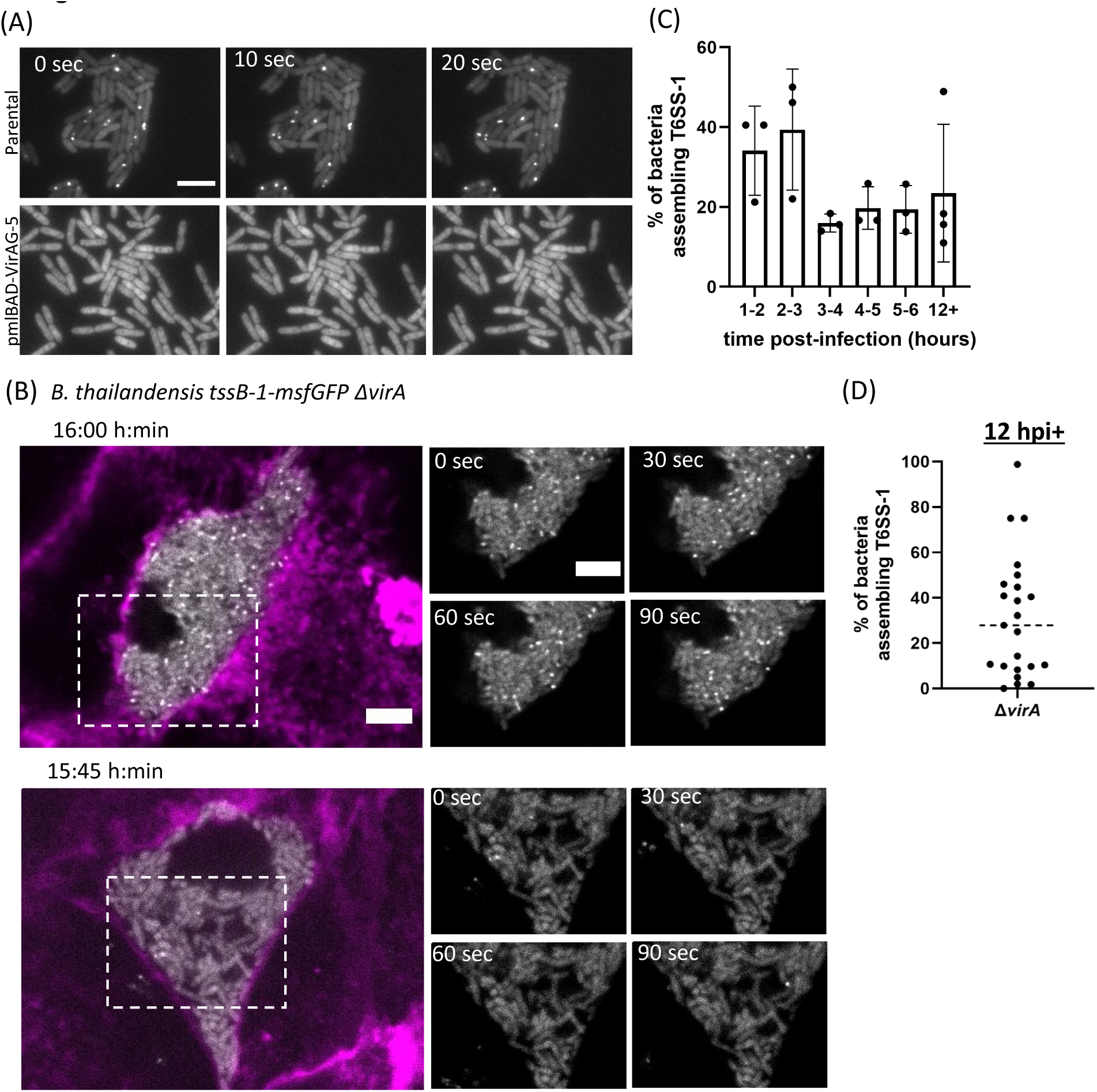
Inhibition of T6SS-1 assemblies by VirAG *in vitro* and during infection. **(A)** Time-lapse microscopy of *B. thailandensis* TssB-1-msfGFP parental strain (top) and a stain harbouring the arabinose-inducible plasmid pmlBAD-VirAG (bottom) on agar pads. Overnight bacterial cultures were subcultured to mid-log phase with 0.4% L-arabinose in LB and imaged on LB agar pads. Scale bar represents 5 µm. **(B)** Two examples of A549 cells infected with *B. thailandensis* Δ*virA*, TssB-1-msfGFP at MOI of 50 at 16 and 15:45 hpi. Host cell membranes were stained with CellMask DeepRed (magenta). Scale bars represent 5 µm. **(C)** Quantification of the percentage of *B. thailandensis* Δ*virA* bacteria that assembled T6SS-1 at different infection time points. A549 cells were infected at an MOI of 200, and at least 100 intracellular bacteria were analyzed per biological replicate (n = 3). Each bacterium was imaged only once in a unique field of view, with time-lapse images acquired every 30 seconds over a 5-minute period. **(D)** Scatter plot showing of the percentage of *B. thailandensis* Δ*virA* assembling T6SS-1 per host cell. A549 cells were infected at MOI 50 and between 12-16 hpi, time-lapse images were acquired every 30 seconds over a 5-minute period. Each data point represents a single host cell, where the total number of bacteria and the number of bacteria assembling T6SS-1 were counted. Data pooled from three biological replicates.

To test this during infection, we constructed a *virA* knock-out strain. To ensure that the T6SS-1 assemblies of the Δ*virA* mutant are functional, we performed a bacterial competition assay with *E. coli* as prey (Fig S6E). Whereas the T6SS-1 deletion strain (Δ*hcp-1*) showed no killing, the WT and Δ*virA* mutant showed similar killing efficiency of *E. coli*, thus confirming that the T6SS-1 assemblies of the Δ*virA* mutant are bactericidal.

We then infected A549 cells with the Δ*virA* mutant and imaged at 16 hpi. Deletion of *virA* allowed *B. thailandensis* to assemble T6SS-1 during infection (Fig 4B, Video 6). The Δ*virA* bacteria accumulated inside of single host cells and failed to spread. This is presumably because the protein required for actin tail motility, BimA, was reported to be not expressed in the absence of VirAG^45^. We then explored if *B. thailandensis* Δ*virA* assembles T6SS-1 at different time points during infection. We infected A549 cells and acquired time-lapse images of 1,834 different bacteria in unique field of views across different time points in three biological replicates. Images were captured every 30 seconds over a 5-minute period to assess whether the bacteria assembled T6SS-1. We found that bacteria assembled T6SS-1 both when inside of dextran-labelled phagosomes or in the cytoplasm (Fig S6A-D). Throughout the imaging period, 15-40% of *B. thailandensis* Δ*virA* assembled T6SS-1 from 1 hpi to 6 hpi (Fig 4C). Interestingly, we noticed a high variability in the percentage of bacteria assembling T6SS-1 at late infection time points (Fig 4D). We imaged 25 infected cells after 12 hpi, with 1,406 bacteria over three biological replicates, and found that in each infected cell, the percentage of bacteria assembling T6SS-1 varied from 1% to 98%.

In addition, since the Δ*virA* mutant lacks actin-tail motility and accumulated in a host cell at high bacterial cell density, we asked if the T6SS-1 assemblies observed in Figure 4B could be caused by increased bacterial cell-cell contact rather than deletion of *virA*. To prevent bacterial spreading and induce accumulation in host cytoplasm, we infected A549 cells with a *virA* positive, *bimA* knock-out strain expressing TssB-1-msfGFP (Fig S6F). Importantly, in the presence of VirA, even when the bacteria were accumulated inside of one host cell and were in contact with one another, no T6SS-1 assemblies were observed suggesting that mere bacterial cell-cell contact or high number of bacteria in the cytoplasm are insufficient to trigger T6SS-1 assembly.

### T6SS-1 activity induces mitochondrial damage

The anti-bacterial T6SS-1 cluster contains BTH_I2698, which was identified as the homolog of Tle1 lipase (Figure S7A) ^43^. We hypothesize that the secretion of T6SS-1 effector lipases could have cross-kingdom effect on the host cells. This led us to examine mitochondrial integrity as a potential intracellular target of these effectors.

To test if *B. thailandensis* T6SS-1 activity causes mitochondrial damage, we infected A549 cells with the *B. thailandensis* Δ*virA* strain that assembles T6SS-1 during infection. There was no difference in the number of bacteria (colony-forming units) in the infected cells between Δ*virA* and Δ*virA/*Δ*hcp-1* mutant (Figure S7B, C) suggesting that the bacterial burden was comparable. To visualize mitochondria, we stained the host cells with Tetramethylrhodamine methyl ester (TMRM), a dye that is fluorescent when sequestered by mitochondria with intact membrane potential. A549 cells were infected, stained and images were taken between 9-10 hpi. Host cells treated with 2 μM of carbonyl cyanide 3-chlorophenylhydrazone (CCCP) was used as a positive control for mitochondrial fragmentation (Figure 5A). We classified three phenotypes of mitochondrial status – healthy, fragmented mitochondria, or mitochondria with no membrane potential (Fig 5B). Healthy mitochondria are tubular, fragmented mitochondria appear as discrete dots, and no fluorescence was detected in mitochondria without membrane potential following TMRM staining. Over 100 infected cells were analyzed for the T6SS-1 assembling Δ*virA* mutant, and the non-assembling Δ*virA/*Δ*hcp-1* mutant. For the Δ*virA* mutant, 12 infected cells displayed a normal, filamentous mitochondrial phenotype, 98 cells showed fragmentation, and 9 cells had no mitochondrial potential (Fig 5C). For the Δ*virA/*Δ*hcp-1* mutant, 47 infected cells displayed a normal, filamentous mitochondrial phenotype, 53 cells showed fragmentation, and 3 cells had no mitochondrial potential.

**Figure 5.**
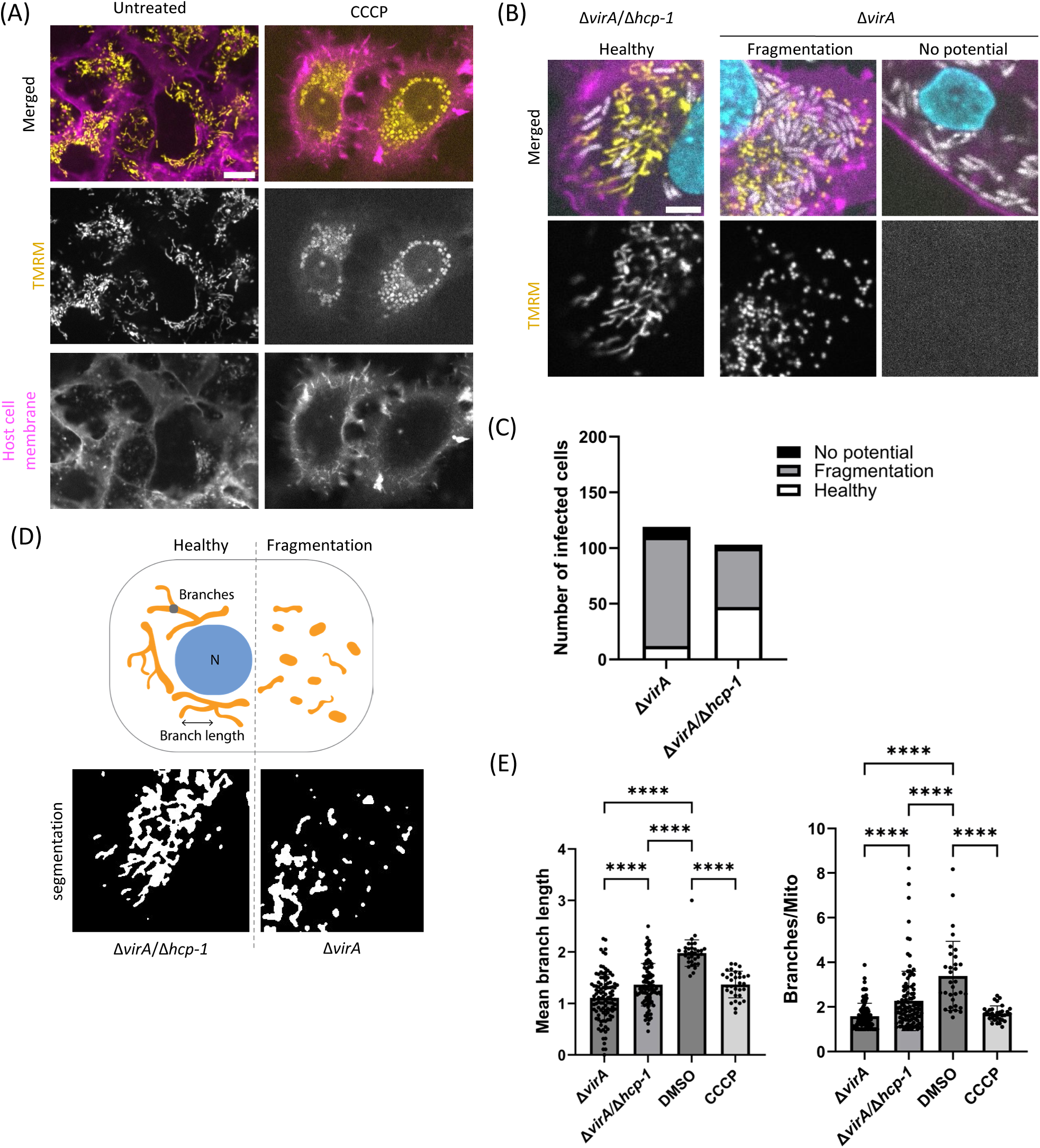
T6SS-1 activity induces mitochondrial damage. **(A)** Tetramethylrhodamine, methyl ester (TMRM) staining of A549 cells under various conditions. A549 cells were either uninfected or treated with 2 µM of CCCP for 30 minutes. Cells were stained with 0.1 µM of TMRM (yellow) and CellMask DeepRed (magenta). Scale bar represents 10 µm. **(B)** Example images of healthy, fragmented mitochondria, and mitochondria with no membrane potential. A549 cells were infected with *B. thailandensis* Δ*virA* or *B. thailandensis* Δ*virA /* Δ*hcp-1* (grey) at MOI 500 for 9-10 hours. Cells were stained with 0.1 µM of TMRM (yellow), CellMask DeepRed (magenta), and Hoechst 33342 (cyan). Scale bar represents 5 µm. **(C)** Proportion of A549 cells infected *B. thailandensis* Δ*virA* or *B. thailandensis* Δ*virA /* Δ*hcp-1* displaying no mitochondrial membrane potential, mitochondrial fragmentation and healthy mitochondrial phenotypes. Data pooled from three biological replicates. **(D)** Schematic representation of healthy and fragmented mitochondrial morphologies illustrating the definitions of “branches per mitochondrion” and “branch length.” N denotes the nucleus. The lower panel shows examples of mitochondria segmented by MitoAnalyzer from the images in (B). **(E)** Automated image analysis of the same dataset from **(A,B)** showing the number of mitochondrial branches per mitochondria and mean mitochondrial branch length performed by the Mitochondria Analyzer plugin for ImageJ. One-way ANOVA was performed and P-values were displayed (****, P < 0.0001).

To quantitatively measure mitochondrial damage, we used the ImageJ plugin Mitochondria Analyzer and quantified the number of branches per mitochondria and the mean mitochondrial branch length^54^. Fragmented mitochondria appear rounder and as isolated puncta, and thus has less branches per mitochondria and shorter branch length. Cells infected with the Δ*virA* mutant showed on average 1.5 branches per mitochondria and with a mean branch length of 1 µm (Fig 5E). Whereas for cells infected with the Δ*virA/*Δ*hcp-1* mutant, they showed on average 2.3 branches per mitochondria and with a mean branch length of 1.4 µm. DMSO-treated uninfected host cells showed on average 3.3 branches per mitochondria and with a mean branch length of 1 µm; whereas CCCP-treated host cells showed on average 1.7 branches per mitochondria and with a mean branch length of 1.3 µm. One-way ANOVA with Tukey’s multiple comparisons test was performed and cells infected with the Δ*virA* mutant had significantly less mitochondrial branching and shorter length, indicating a higher level of mitochondrial fragmentation.

### T6SS-1 activity triggers apoptosis

As mitochondrial fragmentation induces the intrinsic pathway of apoptosis^55,56^, we asked if T6SS-1 activity triggers apoptosis. We used a caspase-3/7 activity detection reagent containing a fluorophore-conjugated peptide substrate specific to caspase-3/7. Upon cleavage by active caspase-3/7, the released fluorophore intercalates with nuclear DNA and emits fluorescence. To optimize the staining conditions, a monolayer of A549 cells were either untreated or treated with staurosphorine, an apoptosis-inducing chemical, were stained with the caspase-3/7 activity detection reagent and apoptotic cells were identified (Fig 6A).

**Figure 6.**
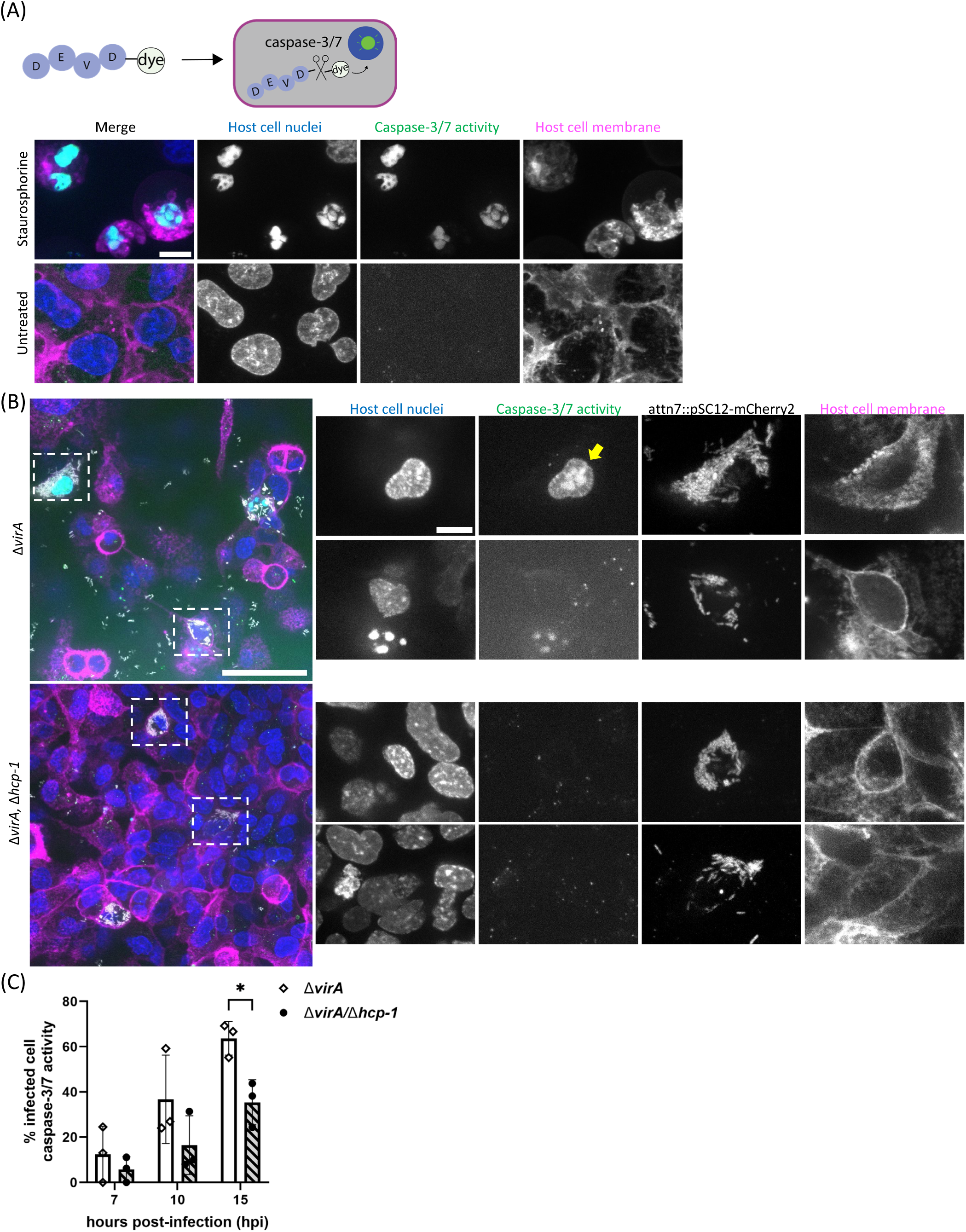
T6SS-1 activity triggers apoptotic cell death. **(A)** Scheme showing the working principle of CellEvent™ Caspase-3/7 Detection Reagent staining and example images of A549 cells that are untreated or treated with staurosporine (1 µM, 18 hours) stained with CellEvent™ Caspase-3/7 Detection Reagent (green). Cells were also stained with Hoechst 33342 (blue) and CellMask DeepRed (magenta). Scale bar represents 10 µm. **(B)** Examples of A549 cells stained with CellEvent™ Caspase-3/7 Detection Reagent (green) and infected with *B. thailandensis* Δ*virA* or *B. thailandensis* Δ*virA /* Δ*hcp-1* (grey) at MOI 500 for 15 hours. A549 cells were also stained with Hoechst 33342 (blue) and CellMask DeepRed (magenta). Yellow arrow indicates host cell nuclei with active caspase-3/7 activity. Scale bars represent 50 µm in the overview image and 10 µm in the magnified image. **(C)** Quantification of the percentage of infected A549 cells displaying caspase-3/7 activity. Cells were infected with either *B. thailandensis* Δ*virA* or *B. thailandensis* Δ*virA /* Δ*hcp-1* (grey) at MOI 500 for 7, 10, or 15 hpi. The total number of infected cells and the number of infected cell nuclei positive for CellEvent™ Caspase-3/7 Detection Reagent were counted. An ordinary two-way ANOVA with Bonferroni’s multiple comparisons test was performed (* P < 0.05).

A549 cells were infected with either the Δ*virA* mutant or the Δ*virA/*Δ*hcp-1* mutant, stained with caspase-3/7 activity detection reagent at 7, 10, 15 hpi, and images were acquired to quantify the number of apoptotic infected cells. At least 20 fields of view (302.28 x 302.28μm) were acquired in each biological replicate. Figure 6B shows two fields of view of infected A549 cells stained with caspase-3/7 activity detection reagent. Host cell nuclei were identified by Hoechst 33342 staining, and a colocalization of the caspase-3/7 activity detection reagent signal with the nuclei was used to identify apoptotic cells. At 7 hpi and 10 hpi, there were approximately twice as many infected cells with active caspase-3/7 when infected with Δ*virA* mutant than when the cells were infected with the the Δ*virA/*Δ*hcp-1* mutant in each biological replicate (Fig 6C). Although some variability was observed in the percentage of apoptotic infected cells, by 15 hpi, 63% of the cells infected the Δ*virA* mutant are apoptotic, while only 35% of those infected with the Δ*virA/*Δ*hcp-1* mutant are apoptotic. This difference was statistically significant. To ensure that the observed difference in cell death was not due to differences in intracellular bacterial burden, we quantified the number of bacteria per infected cell from microscopy images and measured CFU. No differences were observed between the mutants (Figure S7B,C).

## Discussion

The T6SS-1 and T6SS-5 play different roles during *B. thailandensis* lifestyle. Our study showed that the histidine kinase VirAG system in *Burkholderia* spp. inhibits T6SS-1 assemblies during infection and thus prevents T6SS-1 mediated increase in apoptosis via mitochondrial damage.

The lack of intracellular expression of T6SS-2, -4 and -6 is consistent with their unclear role in infection. Previous studies on T6SS-2 and T6SS-6 focused on mice infections and identified no role of the two T6SS in virulence^16^. T6SS-4 is reported active under oxidative stress *in vitro*^57,58^. The anti-eukaryotic T6SS-5’s role in infection has been widely studied in literature^16,17,20,20,41^, however, the temporal dynamics of T6SS-5 expression in live cells remain uncharacterised. By live-cell imaging, we tracked individual bacteria from early stages of infection and identified the time points at which T6SS-5 expression was initiated. Moreover, the anti-bacterial T6SS-1 has primarily been studied in the context of bacterial competition assays.

Here, we identified VirAG as a switch between the T6SS-1 and T6SS-5. The VirAG dimer was reported to be reduced to its active monomer form upon exposure to host cytosolic glutathione (GSH), leading to activation of T6SS-5 expression^47^. In our study, we observed T6SS-5 expression occurring also in late phagosomes before the bacteria appear in the host cytoplasm. Interestingly, we observed T6SS-5 assemblies inside late phagosomes. Although rare, this suggests that the other T6SS-5 components are also expressed at this stage, enabling the formation of functional assemblies. It is plausible that acidification and redox changes accompanying phagosome maturation are sufficient to initiate T6SS activation, which is then accelerated after bacterial escape to the cytosol. Ultimately, both studies support the same overarching conclusion that VirAG is activated and T6SS-5 is expressed during infection.

We show that phagosomal maturation is important for T6SS-5 expression and is consistent with previous studies. Chua *et al.* demonstrated that pH alkalinisation suppresses the anti-eukaryotic T6SS-5 of *B. pseudomallei, B. mallei,* and *B. thailandensis*, as evidenced by the lack of MNGC formation^59^. As MNGC formation is a hallmark for T6SS-5 activity in *Burkholderia*, these findings are consistent with our observation that inhibition of phagosomal acidification traps *B. thailandensis* inside vacuoles and greatly reduce T6SS-5 expression. The rare exceptions that we observed, in which *B. thailandensis* expressed T6SS-5 in bafilomycin-treated cells, may reflect incomplete inhibition of phagosomal maturation in individual host cells, or instances in which cytosolic signals leaked into the phagosome, thereby inducing T6SS-5 expression. Moreover, we concluded that *B. thailandensis* expresses T6SS-5 between 2-3 hpi, and the bacteria mostly are located in Rab7 and LAMP-1 positive late phagosomes. A study by Hu *et al.* reported that between 2-4 hpi, *B. pseudomallei* co-localizes to Rab7 and LAMP-1 positive phagosomes^60^. This suggests that *B. thailandensis* and *B. pseudomallei* likely share the similar intracellular trafficking dynamics.

The anti-bacterial T6SS-1 was shown to assemble in *in vitro* ^44^. Surprisingly, we find that the anti-bacterial T6SS-1 is expressed but not assembled during infection. The gradual decrease in T6SS-1 assemblies matching increase in expression of the anti-eukaryotic T6SS-5 indicates a switch between the two T6SS. It is unclear why T6SS-1 expression would be maintained during infection. One possible explanation is that *B. thailandensis* maintains a pool of T6SS-1 proteins ready to assemble once it exits the host and re-enters the environment. This suggests that the suppression of T6SS-1 inside host cells may be a temporary, reversible adaptation. Interestingly, *B. mallei* has undergone significant genome reduction, and is believed to be evolving towards a more specialized, intracellular lifestyle^61,62^. Consistent with this shift, the anti-bacterial T6SS-1 of *B. mallei* is truncated and likely non-functional^40^. This hints that T6SS-1 is selected against during the evolution of *B. mallei* in the intracellular environment. Hence, we can speculate that *B. thailandensis* inhibits T6SS-1 assemblies during infection as an adaptation to surviving an intracellular environment.

The VgrGs and effectors associated with the anti-bacterial T6SS-1 were identified in three gene clusters located outside the core T6SS locus, each situated at distinct chromosomal locations^43^. Intriguingly, BTH_I2698 is a homolog of Tle1, a lipase in *Pseudomonas aeruginosa*^63^. Moreover, we found that all T6SS-1 VgrGs contain the DUF2345 domain (Fig S7A). This domain is shown to be present in the C-terminal of VgrGs from other bacteria such as VgrG2b from *Pseudomonas aeruginosa*, VgrGi of a clinical strain of *Acinetobacter baumannii*, and VgrG4 from *Klebsiella pneumonia*, all shown to contribute to virulence^64–66^. These observations prompted us to explore the effects of T6SS-1 assemblies to eukaryotic cells. We showed that the Δ*virA* mutant that constitutively assembles T6SS-1 during infection causes more mitochondrial damage and apoptotic cell death than the Δ*virA/*Δ*hcp-1* mutant with inactivated T6SS-1. This was not due to a difference in bacterial load in cells infected with the Δ*virA* mutant since image-based measurements and colony-forming unit analyses showed no significant differences (Figure S7B,C). Our findings suggest that T6SS-1 assemblies induce mitochondrial damage, potentially triggering apoptosis. It is therefore possible that *B. thailandensis* evolved an ability to inhibit T6SS-1 to prevent triggering host cell death and thus preserve its replicative niche.

Overall, our findings demonstrate that VirAG acts as a switch between the anti-eukaryotic T6SS-5 and anti-bacterial T6SS-1 of *B. thailandensis*, enabling the bacterium to better adapt when transitioning from being in the environment to being intracellular. This highlights a regulatory mechanism that governs the utilization of distinct T6SS systems in response to environmental cues.

## Materials and Methods

### Bacterial strains, plasmids and culturing conditions

Bacterial strains used in this study are listed in Table 1. Bacterial strains were grown in LB media overnight at 37 °C with aeration. When appropriate, antibiotics were added at the following concentrations: 50 μg/mL of trimethoprim or 30 μg/mL of gentamicin for *E. coli*; and 200 μg/mL of trimethoprim or 300 μg/mL of kanamycin for *B. thailandensis*.

### Construction of genetic mutants and reporters

Genetic mutants were constructed via allelic exchange via the vector pDONRPEX-18Tp-SceI-pheS or pTOX^67,68^. The homologous flanking regions were at least 1000 bp. pDONRPEX-18Tp-SceI-pheS or pTOX derivatives were conjugated into *B. thailandensis* strains by *E. coli* SM10 λpir. For pDONRPEX-18Tp-SceI-pheS, positive conjugants were selected by trimethoprim resistance and counter selection were performed on M9 minimal medium agar plates supplemented with 0.4% glucose and 0.1% (w/v) 4-chloro-phenylalanine. For pTOX derivatives, all steps were performed in the presence of 2% glucose to prevent premature toxin induction. Conjugants were selected by chloramphenicol resistance and counter selection were performed on M9 minimal medium agar plates supplemented with 2% (w/v) rhamnose.

Bacterial strains constitutively expressing cytoplasmic fluorophores and reporter strains were constructed by chromosomal Tn7 transposase insertion^69^. For constitutive expression, P*_S12_* promoter (*Burkholderia pseudomallei rpsL* promoter) was employed. pUC18tminiTN7Tp and the helper plasmid pTNS2 were conjugated into *B. thailandensis* strains by *E. coli* SM10 λpir and conjugants were selected by trimethoprim resistance. Successful chromosomal insertion into *aat*Tn*7* site downstream of the *glmS* gene were screened by PCR.

### Mammalian cell lines and culturing conditions

The cell lines HeLa cells (CCL-2, ATCC), A549 lung cells (CCL-185, ATCC) were used in this study. HeLa cell lines stably expressing Rab 5 and Rab 7-mApple were provided by Prof. Dr. Anne Spang^70^. The HeLa cells were cultured in high-glucose DMEM with L-glutamine (Sigma-Aldrich) supplemented with 10% heat-inactivated fetal calf serum (FCS, BioConcept AG); and A549 lung cells with Ham’s F-12 media with GlutaMAX™ (Sigma-Aldrich) supplemented with 10% heat-inactivated FCS at 37°C with 5% CO_2_. All experiments were performed between passages 2 to 10 and the cell lines were tested negative for mycoplasma contamination by MycoStrip (InvivoGen).

### Infection assays

Infection assays were performed as described previously^18^. In brief, A549 and HeLa cells were seeded at 8 x 10^4^ cells per well on μ-slide 8 well glass bottom (Ibidi GmbH) 24 hours prior to infection. Bacterial strains were grown overnight in low-salt LB at 37°C with aeration. On the day of the infection assay, overnight bacterial cultures were subcultured in a 1:100 dilution and grown to mid-log phase in low-salt LB. Both the bacterial and mammalian cells were washed twice in Opti-MEM™ Reduced Serum Medium with GlutaMAX™ supplement (Sigma-Aldrich). Bacterial cells were added to the mammalian cells at the specified MOI and the μ-slide 8 well glass bottom were spun down at 300 x g for 5 min. After 1 hour post-infection, Opti-MEM™ supplemented with 300 μg/mL of kanamycin was added to kill extracellular bacteria.

### Immunofluorescence

Cells were infected as described above. For labelling LAMP1, cells were fixed with ice cold 100% methanol for 15 minutes at specified timepoints. The fixed cells were then washed twice with PBS. LAMP1 mouse monoclonal antibody (Abcam) was diluted to 1 μg/mL in 1% Bovine Serum Albumin (BSA, Sigma-Aldrich) and added to the fixed cells to incubate at room temperature for 1 hour. Cells were then washed twice with PBS. Anti-mouse (Abcam) secondary antibodies were diluted 1:300 in 1% BSA and incubated with the cells overnight at 4 degrees. Cells were then washed twice with PBS and imaged.

### Fluorescence microscopy

Infection imaging was performed using a Nikon ECLIPSE Ti2 inverted microscope equipped with a CrestOptics X-Light V3 spinning disk confocal system and a Kinetix sCMOS camera. Images were acquired using a 60× oil immersion objective (NA = 1.42; field of view: 302.28 × 302.28 µm) under live-cell conditions (37 °C, 5% CO₂). Unless otherwise stated, eukaryotic cells were stained with CellMask DeepRed (5 µg/mL) and Hoechst 33342 (40 µM) for 5 min at 37°C with 5% CO_2._

For imaging agar pad, images were acquired on a Nikon Ti-E inverted motorized microscope with Plan Apo 100x Oil Ph3 DM (NA=1.4; field of view – 122.64 x 122.88 μm) objective lens.

### Fluorescence intensity quantification

Time-lapse single plane images were acquired using the confocal microscopy setup described above. Bacteria were segmented with the Otsu thresholding algorithm in ImageJ. The segmented cells were saved as Regions Of Interest (ROIs), and the average fluorescence intensity within each ROI was measured for the relevant fluorescence channels. To account for background signals, fluorescence intensity was also measured from randomly selected ROIs in cell-free regions of the same channel.

### Quantification of T6SS-1 assemblies

Time-lapse imaging was performed using the confocal microscopy setup described above, with images acquired every 30 seconds over a 5-minute period. At each timepoint, a z-stack of three optical sections was captured with a step size of 0.5 µm. Each bacterium was imaged only once in a unique field of view to avoid duplicate counts across timepoints. The images were subsequently processed by maximum intensity projection for analysis. The total number of intracellular bacteria per field was manually counted, and cells assembling T6SS-1 were identified and quantified manually. At least 200 bacteria were counted in each of the three biological replicates.

### Mitochondrial damage quantification

Mitochondria were stained with Tetramethylrhodamine, Methyl Ester, Perchlorate (TMRM, Thermo Fischer) at 100 nM for 15 minutes at 37°C with 5% CO_2_. For positive control, eukaryotic cells were treated with 2 µM of carbonyl cyanide m-chlorophenyl hydrazone (CCCP, Sigma Aldrich) for 30 minutes. Mitochondrial morphology was quantified using the ImageJ plugin Mitochondria Analyzer (https://github.com/AhsenChaudhry/Mitochondria-Analyzer). At least 100 images per experimental condition were acquired across three biological replicates. Infected cells were manually segmented to isolate individual cells for analysis. The mitochondria were thresholded and analyzed with Mitochondria Analyzer. The number of branches per mitochondria and mean branch length were plotted for comparative analysis.

### Cell death assays

Apoptotic cell death were quantified by caspase-3/7 activity (ThermoFischer). Cells were first stained with CellMask DeepRed (5 µg/mL) and Hoechst 33342 (40 µM) for 5 min at 37°C with 5% CO_2_, and washed twice with optimum containing 300 µg/mL kanamycin. CellEvent™ Caspase-3/7 green detection reagent were diluted 1:1000 from a stock concentration of 2 mM to a final working concentration of 2 µM, added directly to the same, and incubated for 30 minutes at 37°C. The cells were imaged without additional washing. Green fluorescence was measured with the 470nm/510nm filter. As a positive control for apoptosis, cells were treated with 1 uM of staurosporine for 18 hours.

### Bacterial competition assay

For quantitative killing assays, mid-log phase bacteria were adjusted to an optical density at 600nm (OD_600_) of 5. Predator and prey species were mixed at 1:1 ratio, and 5 µL of the mixture was spotted on dry LB agar plates and incubated at 30°C for 5 hours. After incubation, the spots were harvested and serially diluted for plating on selective recovery plates (chloramphenicol 150 μg/mL for *E. coli*). CFU was counted after ∼18 hours of incubation at 37°C. Three independent biological replicates were analyzed.

### Image and statistical analysis

The images were analysed with ImageJ/Fiji and GraphPad Prism 9.1.0. were used for data analysis. Two-tailed t-test was performed when comparing two means. Ordinary one-way or two-way ANOVA with Tukey’s or Bonferroni’s multiple comparison test were performed when comparing multiple means. Statistical significance was defined at P<0.05 and the specific statistical testing for each experiment is listed in the figure legends.

## Acknowledgments

The authors would like to thank Prof. Dr. Anne Spang for providing the HeLa cell lines stably expressing Rab 5 and Rab 7-mApple. This work was supported by the Boehringer Ingelheim Fonds PhD fellowship, the University of Basel, and the European Research Council, consolidator grant 865105 – “AimingT6SS”.

## Author contributions

Conceptualization, H.C.C., M.T.W.P., and M.B.; Methodology, H.C.C. and D.J.W.; Investigation, H.C.C., D.J.W., and P.R.I.; Writing – original draft, H.C.C.; writing – review and editing, H.C.C. and M.B.; funding acquisition, M.B.; supervision, M.B.

---

**Summary figure.**
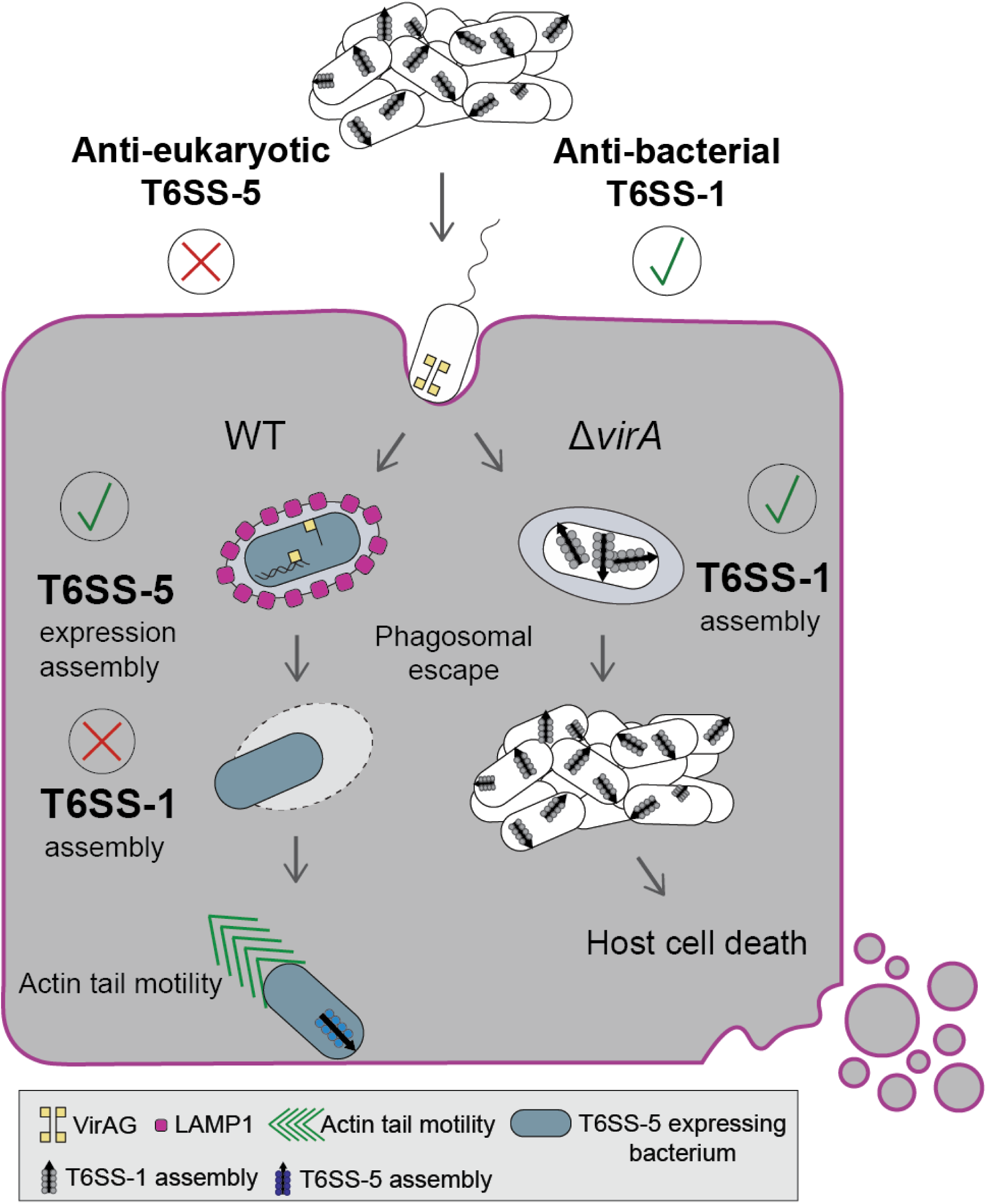

## Supplementary figure legends

**Figure S1.**
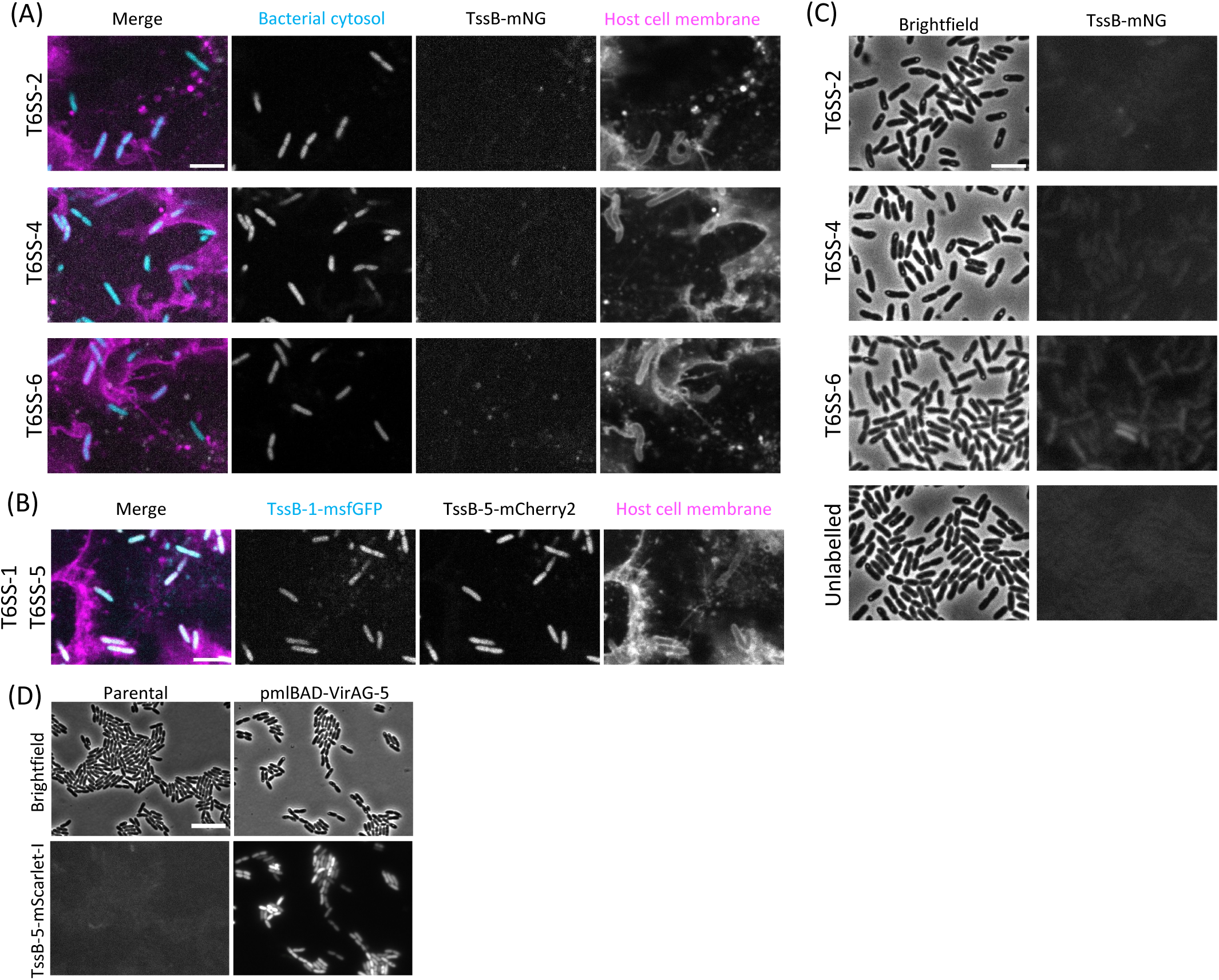
Expression of *B. thailandensis* T6SS. **(A,B)** Representative confocal fluorescence images of *B. thailandensis* E264 infection in A549 cells. Host cell membranes are visualised with CellMask deep red stain (magenta). Images were taken between 11 to 12 hpi and representative of three biological replicates. **(A)** The sheath protein TssB of T6SS-2, T6SS-4 and T6SS-6 were fused with mNeonGreen (mNG) and were shown in grey. The fluorophore mCherry2 was expressed under a ribosomal pSC12 promoter at a mini-Tn7 insertion site on the bacterial chromosome for visualisation and localisation of bacteria in confocal microscopy (cyan). **(B)** TssB-1-msfGFP was shown in cyan and TssB-5-mScarlet-I was shown in grey. **(C)** Wide-field agar pad microscopy of *B. thailandensis* E264 strains shown in (A). Overnight bacterial cultures were subcultured to mid-log phase in LB, adjusted to OD 10, and spotted on LB agar pads. The sheath protein TssB of T6SS-2, T6SS-4 and T6SS-6 were fused with mNeonGreen (mNG) and were shown in grey. Images were adjusted to the same contrast settings. Scale bar represents 5 µm. **(D)** Wide-field agar pad microscopy of parental *B. thailandensis* with chromosomal TssB-5-mScarlet-I and *B. thailandensis* with chromosomal TssB-5-mScarlet-I harbouring the arabinose-inducible plasmid pmlBAD-VirAG. Overnight bacterial cultures were subcultured to mid-log phase in LB supplemented with 0.4% L-arabinose. The cultures were adjusted to OD 10 and spotted on LB agar pads. Scale bar represents 10 µm.

**Figure S2.**
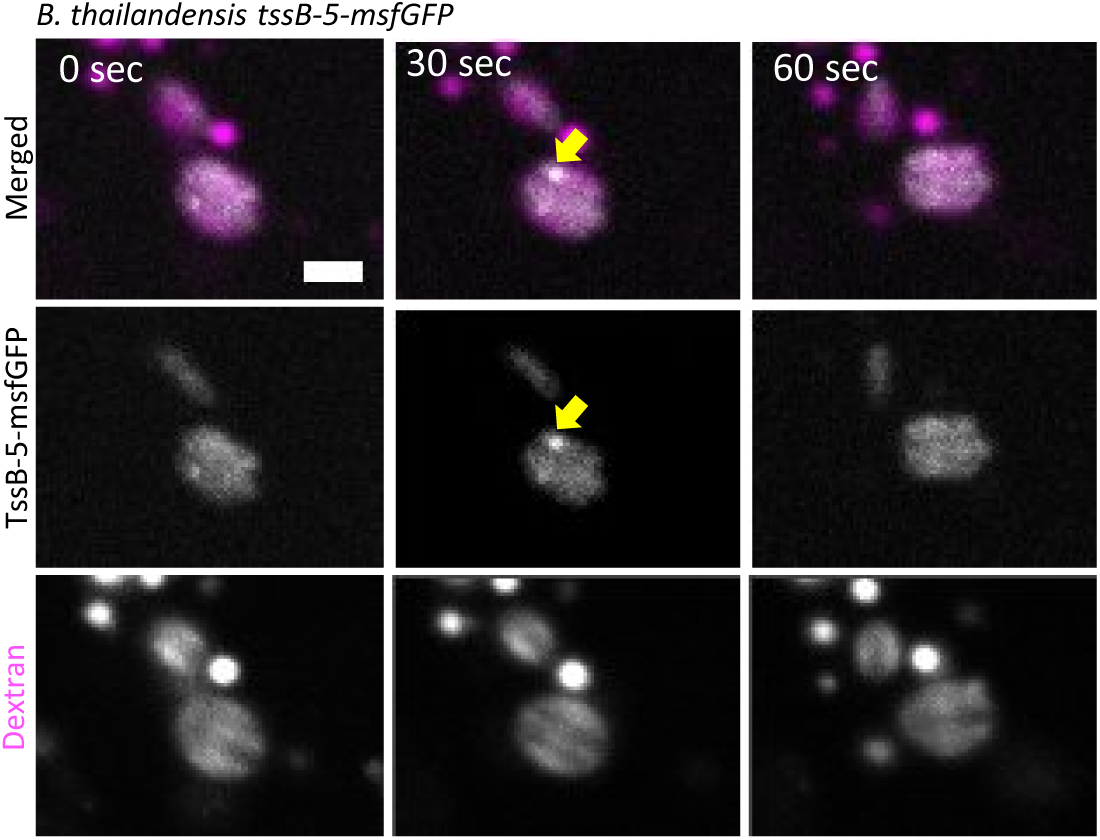
T6SS-5 assemblies in phagosome. *B. thailandensis* assembling T6SS-5 at 3 hpi inside of a dextran-labelled vacuole during infection of A549 cells. A549 cells were pre-treated with 0.25 mg/mL of fluorescent dextran to identify bacteria inside phagosomes. Yellow arrows indicate a T6SS-5 assembly event. Scale bar represents 2 µm.

**Figure S3.**
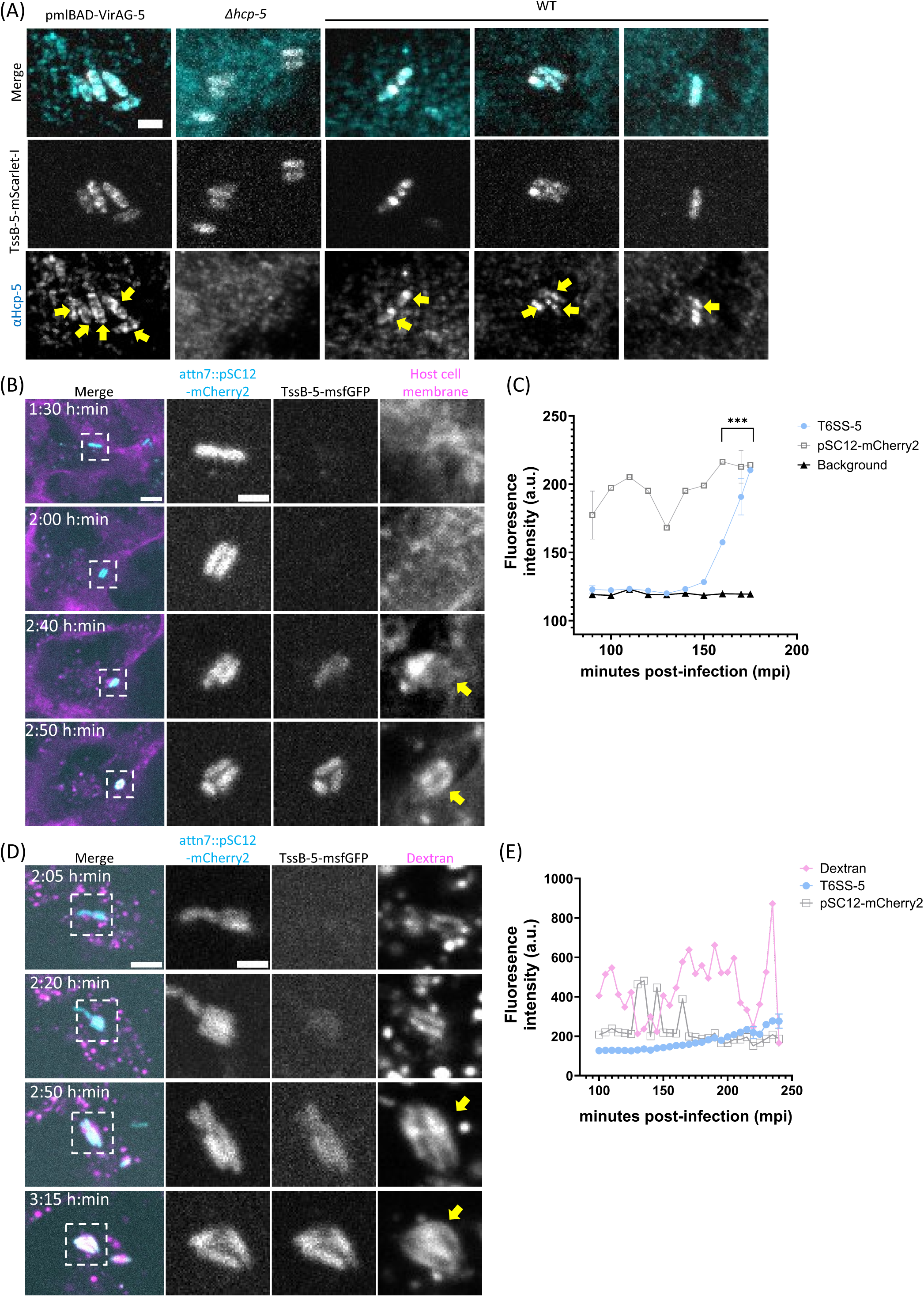
Expression of the anti-eukaryotic T6SS-5 structural and regulatory components *in vitro* and during infection. **(A)** Immunofluorescence staining of Hcp-5. A549 cells were infected with various *B. thailandensis* strains at MOI 500 and fixed with 4% PFA at 3 hpi. The slide was then permeabilized with lysozyme and mutanolysin for 1 hour at 37 degrees. Then stained with anti-hcp-5 antibody at 1 ug/mL for 1 hour at room temperature. Then the slide was washed and stained with anti-rabbit-AlexaFluor488 overnight at 4 degrees. Scale bar represents 2 µm. Yellow arrows indicate Hcp-5 expression. **(B)** Time-lapse imaging of A549 cells infected with *B. thailandensis* expressing cytosolic mCherry2 under ribosomal promoter pSC12 (cyan) and TssB-5-msfGFP (grey) at MOI of 50. Host cell membrane was stained with CellMask DeepRed (magenta). Yellow arrows indicate host cell membrane around the bacteria. Scale bars represent 5 µm in the overview image and 5 µm in the magnified image. **(C)** Fluorescence intensity of various channels during the infection period shown in (B). Bacteria were segmented and tracked, and fluorescence intensities of TssB-5-msfGFP (T6SS-5, in blue), bacterial cytosol (pSC12-mCherry2, in grey), and background (black) were plotted. A two-way ANOVA with Tukey’s multiple comparisons test was performed (*** P<0.0001). **(D)** Time-lapse imaging of A549 cells infected with *B. thailandensis* expressing cytosolic mCherry2 under ribosomal promoter pSC12 (cyan) and TssB-5-msfGFP (grey) at MOI of 200. Host cells were pre-treated with 0.25 mg/mL of fluorescent dextran for 18 hours (magenta). Yellow arrows indicate bacteria expressing T6SS-5 when inside of a dextran-labelled vacuole. Scale bars represent 5 µm in the overview image and 5 µm in the magnified image. **(E)** Fluorescence intensity of various channels during the infection period shown in (D). Bacteria were segmented and tracked, and fluorescence intensities of TssB-5-msfGFP (T6SS-5, in blue), bacterial cytosol (pSC12-mCherry2, in grey), and fluorescent dexran (magenta) were plotted.

**Figure S4.**
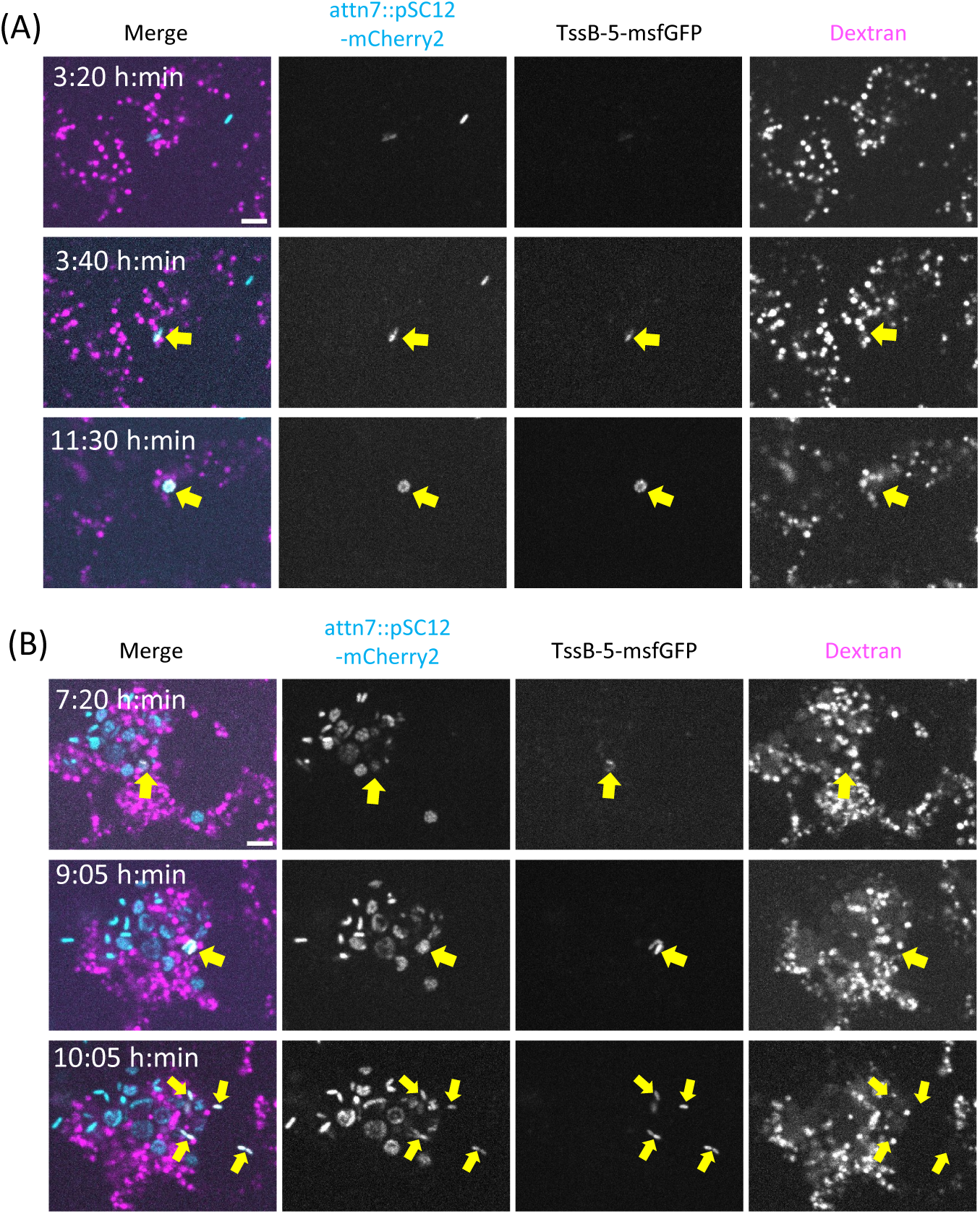
Representative examples of T6SS-5 expression upon bafilomycin A1 treatment. Examples of bacterial clusters that **(A)** remain within the phagosome and **(B)** escape from the phagosome. Time-lapse imaging of A549 cells treated with bafilomycin A1 (100nM) or DMSO infected with *B. thailandensis* expressing cytosolic mCherry2 under ribosomal promoter pSC12 (cyan) and TssB-5-msfGFP (grey) at MOI of 200. A549 cells were pre-treated with 0.25 mg/mL of fluorescent dextran for 18 hours to identify intra-phagosomal bacteria. Host cells were pre-treated 1 hour before infection, infected with *B. thailandensis* for 1 hour without treatment, and bafilomycin A1 or DMSO was added and kept in the media with 300 µg/mL kanamycin from 1 hpi onwards. Host cell membrane was stained with CellMask DeepRed (magenta). Yellow arrows indicate bacteria that express T6SS-5. Scale bars represent 5 µm.

**Figure S5.**
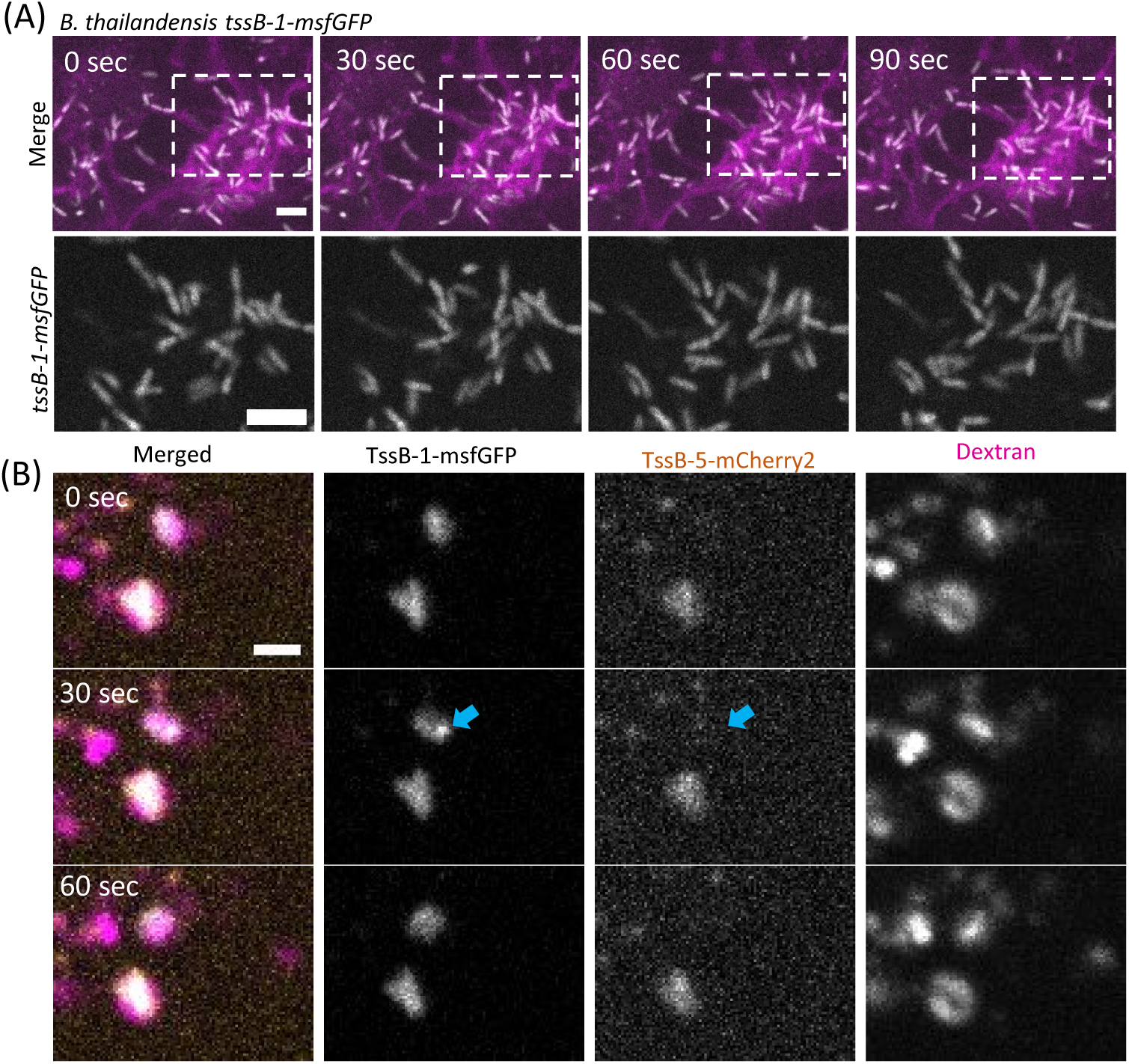
T6SS-1 assembly dynamics during infection. **(A)** Time-lapse microscopy of A549 cells infected with *B. thailandensis* TssB-1-msfGFP at MOI of 50 at 13 hpi. Host cell membranes were stained with CellMask DeepRed (magenta). Scale bars represent 5 µm. **(B)** Time-lapse microscopy of A549 cells infected with *B. thailandensis* expressing TssB-1-msfGFP and TssB-5-mCherry2 at MOI of 500 at 3 hpi. Host cells were pre-treated with 0.25 mg/mL of fluorescent dextran for 18 hours (magenta). Blue arrows indicate a bacteria assembling T6SS-1 and not expressing T6SS-5. Scale bar represents 2 µm.

**Figure S6.**
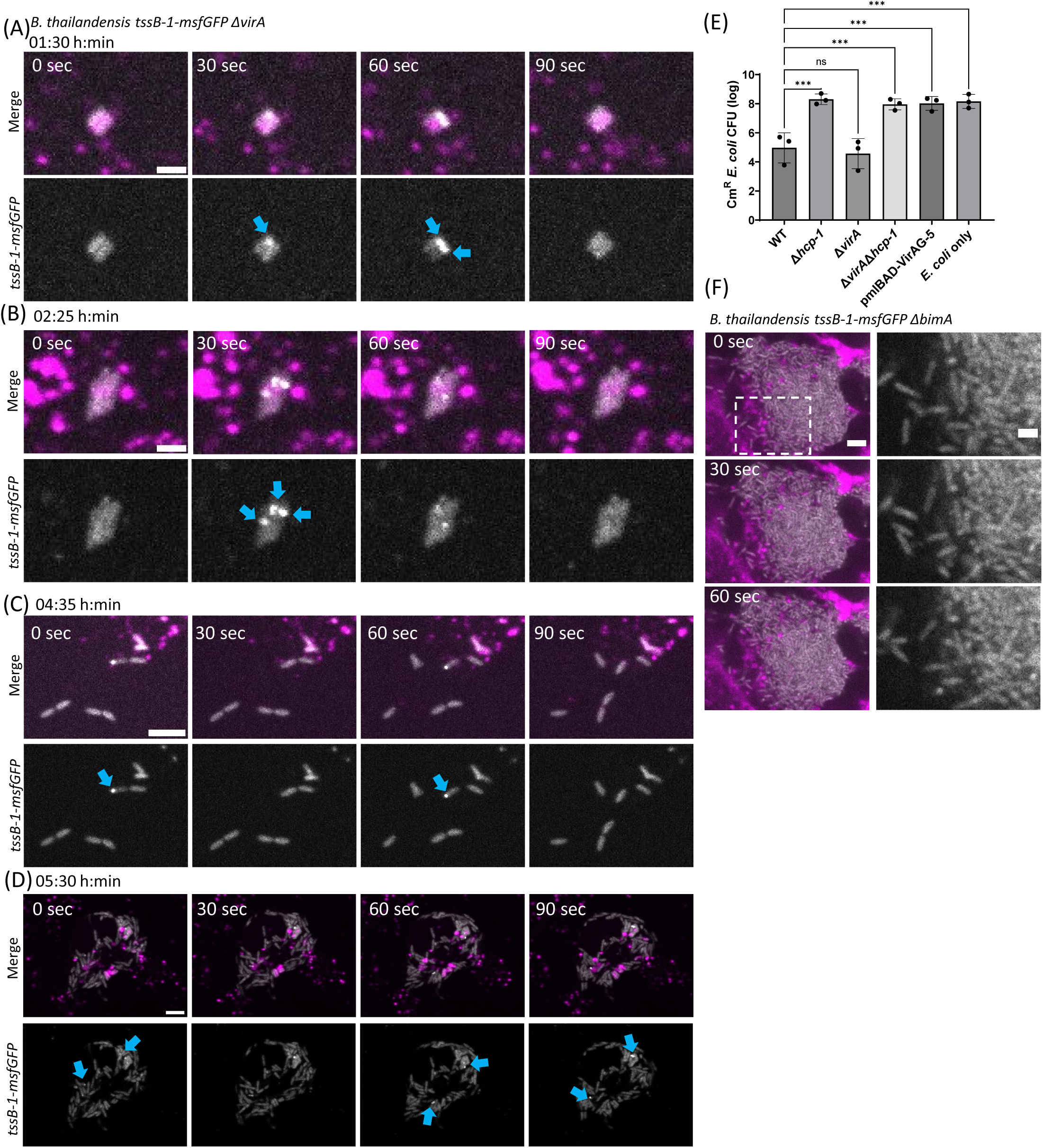
Intracellular T6SS-1 dynamics and bacterial competition assay. Time-lapse microscopy of A549 cells infected with *B. thailandensis ΔvirA,* TssB-1-msfGFP at MOI of 50 at **(A)** 1:30 hpi, **(B)** 2:25 hpi, **(C)** 4:35 hpi, and **(D)** 5:30 hpi. A549 cells were pre-treated with 0.25 mg/mL of fluorescent dextran (magenta) to identify bacteria inside phagosomes. Blue arrows indicate T6SS-1 assembly events. Scale bars represent 2 µm for (A & B) and 5 µm for (C & D). **(E)** Recovery of viable *E. coli* prey after competition with *B. thailandensis* strains. *E. coli* prey harbouring a plasmid conferring trimethoprim resistance was co-cultured with *B. thailandensis* strains at 1:1 ratio on LB agar plates for 5 hours at 30 degrees. The co-culture were harvested and serially diluted and spotted on LB agar plates containing 50 µg/mL trimethoprim. Colony forming units (CFU) of *E. coli* was plotted. Ordinary one-way ANOVA with Dunnett’s multiple comparisons test was performed (*** P< 0.001, ns = not significant). **(F)** Time-lapse microscopy of A549 cells infected with *B. thailandensis* Δ*bimA*, TssB-1-msfGFP at MOI of 50 at 14 hpi. Host cell membranes were stained with CellMask DeepRed (magenta). Scale bars represent 5 µm.

**Figure S7.**
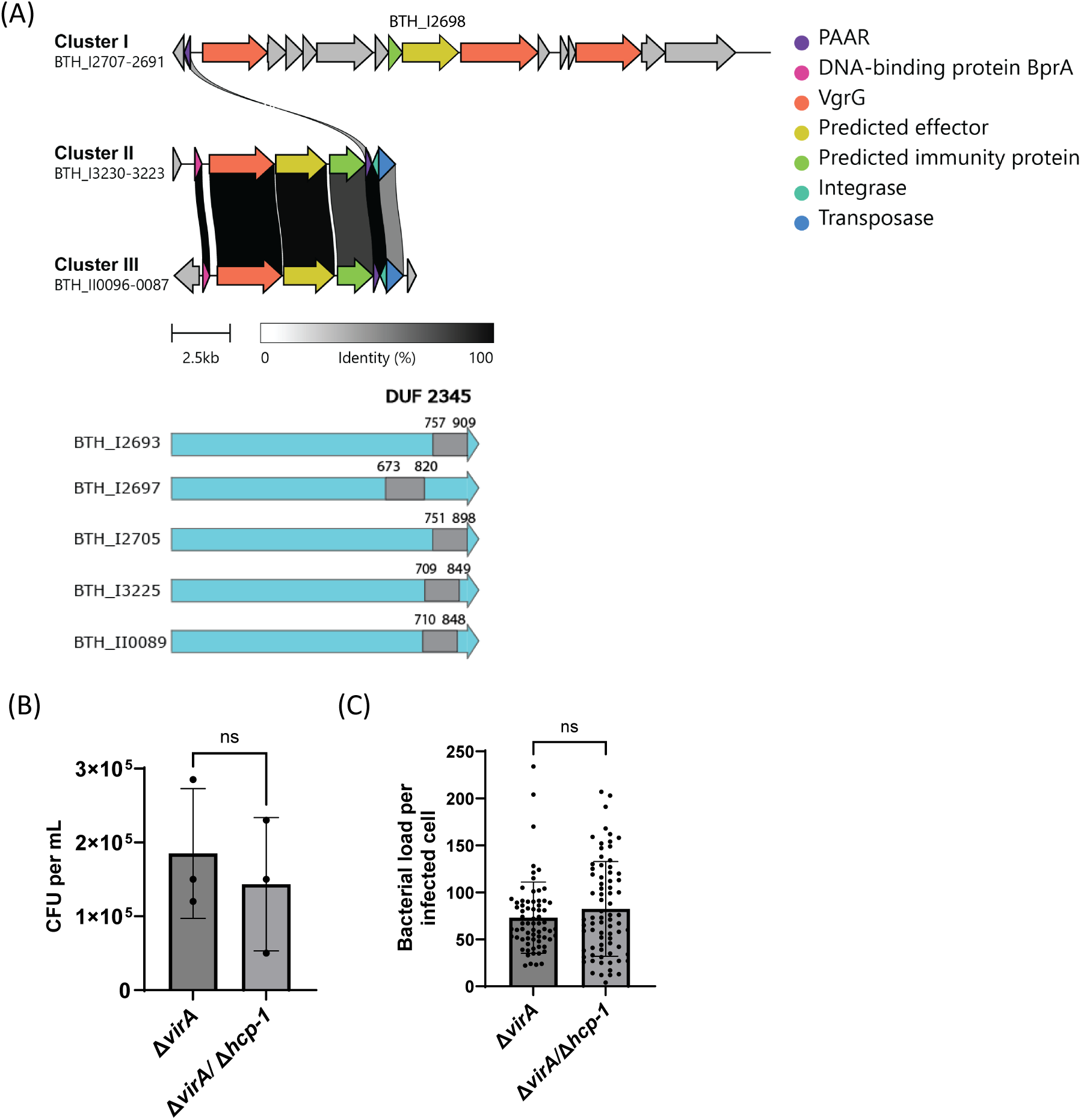
Genomic organization of the T6SS-1 cluster. **(A)** Genomic clusters of the T6SS-1 components. Spike protein VgrGs are colour-coded blue, PAAR motif in green, and transposon elements in red. Synteny was determined by CAGEcat^71^. **(B)** Colony forming units (CFU) and the number of bacteria per A549 cells infected with *B. thailandensis ΔvirA,* or *B. thailandensis* Δ*virA /* Δ*hcp-1*. Infected cells were lysed at 15 hpi with 0.1% triton X-100 and the lysate was serially diluted and plated on LB plate for counting CFU. Number of bacteria per infected cell was manually quantified based on the dataset in Figure 5. Data represents three biological replicates. Unpaired t-test with Welch’s corrections was performed (ns = not significant). **(C)** Number of bacteria per infected cell. A549 cells were infected with either *B. thailandensis ΔvirA,* or *B. thailandensis* Δ*virA /* Δ*hcp-1*. Images were acquired between 9-10 hpi, and the number of bacteria per infected cell were manually counted. Each point represents an infected. Data pooled from three biological replicates. Unpaired t-test with Welch’s corrections was performed (ns = not significant).

## Video legends

**Video 1. T6SS-5 expression during infection.** This video shows two segments (Example 1-2) of T6SS-5 expression inside of A549 cells. Each field of view is 32.50 x 24.38 μm. T6SS-5 is visualized with a TssB-5-mScarlet-I protein fusion and are shown in grayscale. Bacterial cytosol shown in cyan. Host cell membrane is stained with CellMask and shown in magenta. The time-lapse video is shown with 10 frames per second. The scale bars represent 5 μm.

**Video 2. T6SS-5 expression in dextran-labelled phagosomes.** This video shows two segments (Example 1-2) of T6SS-5 expression inside of A549 cells. Each field of view is 32.50 x 24.38 μm. T6SS-5 is visualized with a TssB-5-mScarlet-I protein fusion and are shown in grayscale. Bacterial cytosol shown in cyan. Host cells were pre-treated with 0.25 mg/mL of fluorescent dextran for 18 hours (magenta). The time-lapse video is shown with 10 frames per second. The scale bars represent 5 μm.

**Video 3. Bafilomycin A1 inhibits T6SS-5 expression.** This video shows four segments of *B. thailandensis* infection of A549 cells either treated with 100nM bafilomycin A1 (BAF) or DMSO. Two fields of view (44.45 x 33.34 μm) from independent biological replicates are shown per condition. T6SS-5 is visualized with a TssB-5-mScarlet-I protein fusion and are shown in grayscale. Bacterial cytosol shown in cyan. Host cell membrane is stained with CellMask and shown in magenta. The time-lapse video is shown with 15 frames per second. The scale bars represent 5 μm.

**Video 4. T6SS-1 does not assemble during infection.** This video shows three segments (Examples 1-3) of T6SS-1 dynamics inside of A549 cells at 13 hpi. The three fields of view (43.33 x 32.50 μm) are from independent biological replicates. T6SS-1 is visualized with a TssB-1-msfGFP protein fusion and are shown in grayscale. Host cell membrane is stained with CellMask and shown in magenta. The time-lapse video is shown with 10 frames per second. The scale bars represent 5 μm.

**Video 5. Inhibition of T6SS-1 assemblies by VirAG *in vitro*.** Time-lapse microscopy of *B. thailandensis* TssB-1-msfGFP parental strain (top row) and a stain harbouring the arabinose-inducible plasmid pmlBAD-VirAG (bottom row) on agar pads. Three fields of view (26.00 x 19.50 μm) from independent biological replicates are shown per strain. The time-lapse video is shown with 6 frames per second. The scale bars represent 5 μm.

**Video 6. *B. thailandensis* Δ*virA* assembles T6SS-1 during infection.** This video shows four segments (Examples 1-4) of *B. thailandensis* Δ*virA* T6SS-1 assemblies inside of A549 cells between 13-16 hpi. T6SS-1 is visualized with a TssB-1-msfGFP protein fusion and are shown in grayscale. Host cell membrane is stained with CellMask and shown in magenta. The time-lapse video is shown with 10 frames per second. The scale bars represent 5 μm.

