## Supplemental Table 1 for "Regulation of Type VI secretion systems of *Burkholderia thailandensis* during transition to intracellular lifestyle"

| **Reagent or Resource** | **Source** | **Identifier** |
| --- | --- | --- |
| **Bacterial strains** | | |
| *Burkholderia thailandensis* E264 wild-type | Ku *et al.* ^1^ | N/A |
| *Burkholderia thailandensis Δhcp-5, tssB-5-mScarletI* | Plum *et al.* ^2^ | N/A |
| *Burkholderia thailandensis* tssB-5-mScarlet-I | Plum *et al.* ^2^ | N/A |
| *Burkholderia thailandensis* tssB-5-msfGFP | Plum *et al.* ^2^ | N/A |
| *Burkholderia thailandensis* tssB-1-msfGFP | Lin *et al.* ^3^ | N/A |
| *Burkholderia thailandensis* tssB-1-msfGFP, TssB-5-mCherry2 | This study | N/A |
| *Burkholderia thailandensis* tssB-2-mNeonGreen | This study | N/A |
| *Burkholderia thailandensis* tssB-4-mNeonGreen | This study | N/A |
| *Burkholderia thailandensis* tssB-6-mNeonGreen | This study | N/A |
| *Burkholderia thailandensis* tssB-5-mScarlet-I, pmlBAD-VirAG-5 | This study | N/A |
| *Burkholderia thailandensis BTH_I_TN7L::pCS12-mCherry2, tssB-5-msfGFP* | This study | N/A |
| *Burkholderia thailandensis* E264 Δ*virA* | This study | N/A |
| *Burkholderia thailandensis* E264 Δ*virA* Δ*hcp-1* | This study | N/A |
| *Burkholderia thailandensis* E264 Δ*virA*, tssB-1-msfGFP | This study | N/A |
| *Burkholderia thailandensis* E264 Δ*virA* Δ*hcp-1,* tssB-1-msfGFP | This study | N/A |
| *Burkholderia thailandensis ΔbimA, tssB-1-msfGFP* | This study | N/A |
| **Oligonucleotides** | | |
| TN7L_PC12-mCherry2.FOR | Plum *et al.* ^2^ | TCAGTAGGGCCCTTACTTGTACAGCTCGTCCATGCC |
| TN7L_PC12-mCherry2.REV |  | AGTCATGGTACCAGCTGTTGACTCGCTTGGGATTTTCGGAATATCATGCCGGGTCTGACAATCGGATCGAGCTTCGAAAGGACAAGCATATGGTGAGCAAGGGCGAGGAGGATAAC |
| BTH_II_TN7_Det.FOR |  | GTCGGTGACGGTGGAGTAG |
| BTH_II_TN7_Det.Rev |  | TGTGAATGGTCAGACGCTGTTCG |
| BTH_I_TN7_Det.FOR |  | GCGTTCGTCGTCCACTGGGA |
| BTH_I_TN7_Det.Rev |  | CCGTGACGGTGGAGTAAAG |
| pmlBAD-VirAG-5_KpnI-EcoRI.FOR | This study | TACTGTTTCTCCATACCCGTTTTTTTGGGCTAGCAGGAGGAATTCTTGCTGCTGCTGTTCTACGCG |
| pmlBAD-VirAG-5_KpnI-EcoRI.REV |  | TTGCATGCCTGCAGGTCGACTCTAGAGGATCCCCGGGTACCTCATTCCCATAGCGTCTCCACGT |
| TssB-2-mNG-F1.F |  | CACGACGTTGTAAAACGACGGCCAGTCTTAAGCTCGGGCCCGACACGCATCCGGTTG |
| TssB-2-mNG-F1.R |  | ACTCCTCCTCCTGCGGCCGCGTGCTTTTCCTTCGATTCCC |
| TssB-2-mNG-F2.F |  | GGGAATCGAAGGAAAAGCACGCGGCCGCAGGAG |
| TssB-2-mNG-F2.R |  | CAGTGCTTTTCCTTCGATTCTTTGTACAGCTCATCCATACCC |
| TssB-2-mNG-F3.F |  | GTATGGATGAGCTGTACAAAGAATCGAAGGAAAAGCACTGATTC |
| TssB-2-mNG-F3.R |  | GGAAACAGCTATGACATGATTACGAATTCGAGCTCGGTACCCGGCAGGTTCTCGACGAC |
| TssB-2-mNG-det.1 |  | GTTCTGATCGTCGGGCATTC |
| TssB-2-mNG-det.2 |  | ATCAACCATTGCTGCCTGAAA |
| TssB-2-mNG-det.3 |  | GTTTCTGGAAATCGTCGTGAT |
| TssB-2-mNG-det.4 |  | GTCGGTGTTCTTTCGATGAAT |
| TssB-4-mNeonGreen_F1.FOR |  | CACGACGTTGTAAAACGACGGCCAGTCTTAAGCTCGGGCCCCGTGAGCGCGAGGTAGCG |
| TssB-4-mNeonGreen_F1.REV |  | GTATGGATGAGCTGTACAAACCGGAAGAGGGTGCG |
| TssB-4-mNeonGreen_F2.FOR |  | CATTTCGCACCCTCTTCCGGTTTGTACAGCTCATCCATACC |
| TssB-4-mNeonGreen_F2.REV |  | AACCGGAAGAGGGTGCGAAAGCGGCCGCAGGAG |
| TssB-4-mNeonGreen_F3.FOR |  | ACTCCTCCTCCTGCGGCCGCTTTCGCACCCTCTTCCG |
| TssB-4-mNeonGreen_F3.REV |  | GGAAACAGCTATGACATGATTACGAATTCGAGCTCGGTACCGGATCGTCGAGTACTTCC |
| TssB-4-mNG-det.For |  | CATCGCATAGGCGGAGTTC |
| TssB-4-mNG-det.Rev |  | CTGAGATCGAGGCAGGAAGTC |
| T6SS-6-mNG_F1.FOR |  | CACGACGTTGTAAAACGACGGCCAGTCTTAAGCTCGGGCCCGCGAACGGCCGCG |
| T6SS-6-mNG_F1.REV |  | ACTCCTCCTCCTGCGGCCGCTTCATGGCGAATCTCCGAATCG |
| T6SS-6-mNG_F2.FOR |  | ATTCGGAGATTCGCCATGAAGCGGCCGCAGGAGGAG |
| T6SS-6-mNG_F2.REV |  | TGGCGAATCTCCGAATCGTCTTTGTACAGCTCATCCATACCCATCACATC |
| T6SS-6-mNG_F3.FOR |  | GTATGGATGAGCTGTACAAAGACGATTCGGAGATTCGCC |
| T6SS-6-mNG_F3.REV |  | GGAAACAGCTATGACATGATTACGAATTCGAGCTCGGTACCGAATTCGCCCACGTATAGCGGT |
| TssB-6-mNG-seq.1 |  | ACACGAGCGACTTCGAGA |
| TssB-6-mNG-seq.2 |  | ATGACGTTCGAGAGCATCGAC |
| TssB-6-mNG-seq.3 |  | GATGAACGAACGGCATGA |
| Hcp1-KO-F1.FOR |  | TCAGTAAAGCTTGGCACTGCACGCC |
| Hcp1-KO-F1.REV |  | GCATTTGAAGGACAAGACCTAC |
| Hcp1-KO-F2.FOR |  | GGTCTTGTCCTTCAAATGCAT |
| Hcp1-KO-F2.REV |  | TCAGTATCTAGAAGCAGATGTCGCATGT |
| dHcp1_chromo_det.1 |  | CACTGCACGCCTCGAATC |
| dHcp1_chromo_det.end |  | CTCTTCAAGAAGGTGTACGAAGAA |
| dVirA_F1.F |  | CACGACGTTGTAAAACGACGGCCAGTCTTAAGCTCGGGCCCGCGTGATCGGATACGC |
| dVirA_F1.R |  | CATCGTTTCCTCTCCTGTACCGCGTAGAACAGCAGCAG |
| dVirA_F2.F |  | TGCTGCTGCTGTTCTACGCGGTACAGGAGAGGAAACGATGAATG |
| dVirA_F2.R |  | GGAAACAGCTATGACATGATTACGAATTCGAGCTCGGTACGAGCAGCCATTGCAGCG |
| dVirA_chromo_det_beg |  | CTCGACGAGGAACTTCAGC |
| dVirA_chromo_det_2 |  | GATGTTGACCCGCTCTTTC |
| dVirA_chromo_det.end |  | ATGCTCGGCGAAGGAAAG |
| BimA-KO_F1.FOR |  | CACGACGTTGTAAAACGACGGCCAGTCTTAAGCTCGGGCCCCGCGCGCCGAGC |
| BimA-KO_F1.REV |  | TGTGTTCATCCATCGCACCGGGCATGAGCTGGCAATGG |
| BimA-KO_F2.FOR |  | CACCATTGCCAGCTCATGCCCGGTGCGATGGATGAACAC |
| BimA-KO_F2.REV |  | GGAAACAGCTATGACATGATTACGAATTCGAGCTCGGTACCGGCTGGTTCGGCTATGCG |
| BimA-KO-Det.For |  | CAAATGCCGCGTTGACAT |
| BimA-KO-Det.Rev |  | CATACCGATACGCCGCTGT |
| **Recombinant DNA** | | |
| pDONRPEX18Tp-SceI-pheS | Addgene | Plasmid#11608 |
| pUC18T-mini-Tn7T-Tp | Addgene | Plasmid#65024 |
| pUC18T-mini-Tn7T-Tp-*pSC12-mCherry2* | Plum *et al.* ^2^ | N/A |
| pTNS2 | Addgene | Plasmid#64968 |
| pmlBAD | Chen *et al.*^4^ | N/A |
| pmlBAD-VirAG-5 | This study | N/A |
| pDONRPEX18Tp-TssB-2-mNeonGreen | This study | N/A |
| pDONRPEX18Tp-TssB-4-mNeonGreen | This study | N/A |
| pDONRPEX18Tp-TssB-6-mNeonGreen | This study | N/A |
| pDONRPEX18Tp-Δ*hcp-1* | This study | N/A |
| pDONRPEX18Tp-Δ*virA* | This study | N/A |
| pDONRPEX18Tp-Δ*bimA* | This study | N/A |
| **Cell lines** | | |
| Human: A549 cells | ATCC | Cat# 86012804  Lot# 18F020 |
| Human: HeLa mApple-Rab5 | Podinovskaia *et al.*^5^ |  |
| Human: HeLa mApple-Rab7 | Podinovskaia *et al.*^5^ |  |
